# Index and biological spectrum of accessible DNA elements in the human genome

**DOI:** 10.1101/822510

**Authors:** Wouter Meuleman, Alexander Muratov, Eric Rynes, Jessica Halow, Kristen Lee, Daniel Bates, Morgan Diegel, Douglass Dunn, Fidencio Neri, Athanasios Teodosiadis, Alex Reynolds, Eric Haugen, Jemma Nelson, Audra Johnson, Mark Frerker, Michael Buckley, Richard Sandstrom, Jeff Vierstra, Rajinder Kaul, John Stamatoyannopoulos

## Abstract

DNase I hypersensitive sites (DHSs) are generic markers of regulatory DNA and harbor disease- and phenotypic trait-associated genetic variation. We established high-precision maps of DNase I hypersensitive sites from 733 human biosamples encompassing 439 cell and tissue types and states, and integrated these to precisely delineate and numerically index ~3.6 million DHSs encoded within the human genome, providing a common coordinate system for regulatory DNA. Here we show that the expansive scale of cell and tissue states sampled exposes an unprecedented degree of stereotyped actuation of large sets of elements, signaling the operation of distinct genome-scale regulatory programs. We show further that the complex actuation patterns of individual elements can be captured comprehensively by a simple regulatory vocabulary reflecting their dominant cellular manifestation. This vocabulary, in turn, enables comprehensive and quantitative regulatory annotation of both protein-coding genes and the vast array of well-defined but poorly-characterized non-coding RNA genes. Finally, we show that the combination of high-precision DHSs and regulatory vocabularies markedly concentrate disease- and trait-associated non-coding genetic signals both along the genome and across cellular compartments. Taken together, our results provide a common and extensible coordinate system and vocabulary for human regulatory DNA, and a new global perspective on the architecture of human gene regulation.

## Introduction

A fundamental goal of the ENCODE project is to delineate with the highest possible precision the repertoire of regulatory DNA elements encoded within the human genome sequence. A canonical feature of actuated cis-regulatory elements – promoters, enhancers, silencers, chromatin insulators/enhancer blockers, and locus control regions – is focal alteration in chromatin structure resulting in heightened DNA accessibility to nucleases and other protein factors ^1,2^. From their discovery 40 years ago ^3–5^, DNase I hypersensitive sites (DHSs) have provided reliable signposts for high-precision regulatory DNA delineation in the human and other complex genomes ^1,5–8^. DHSs typically mark compact (<250bp) elements densely populated by sequence-specific regulatory factors; these factors, in turn, produce nucleotide-resolution ‘footprints’ when exposed to DNase I^9^ that further illuminate the underlying regulatory architecture and logic of regulatory DNA^10^.

Following the sequencing of the human genome, the advent of genome-scale mapping of DHSs ^11–14^ and its application to diverse human and mouse cell and tissue types ^15,16^ has yielded many insights into the organization ^15^, evolution ^16–18^, activity ^15,19,20^, and function ^15,19,21^ of human regulatory DNA in both normal and malignant states ^22^. Cell type- and state-selectivity of DNA accessibility is a cardinal property of regulatory DNA, with only a small fraction of all genome-encoded elements becoming actuated in a given cellular context ^15,22^. Most cell-selective behavior is manifested at distal, non-promoter elements, which comprise the plurality of DHSs ^15^. Exceptions to this are the minority of DHSs that mark active or potentiated transcriptional start sites (TSSs), or those arising from non-promoter occupancy sites for CTCF ^23–25^. While it is possible to infer functional properties of a subset of DHSs on the basis of surrounding histone modification patterns ^26–28^, such features do not account well for complex behaviors such as primed elements poised to receive environmental or other stimuli ^19^, or quiescent but developmentally actuated elements ^22^.

The overwhelming majority of disease- and trait-associated variants identified by genome-wide association studies (GWAS) lie in non-coding regions of the genome, and are strongly enriched in DHSs, particularly from disease-relevant cell and tissue types ^29,30^. Additionally, DHSs harbor the subset of GWAS variants that account for the majority of trait heritability explained by genotyped SNPs ^31^. Beyond these general principles, which derive from simple localization of variants within DHSs, deeper insights have been generally limited by the lack of comprehensive DHS annotations that capture their biological behavior.

The above findings have collectively required a combination of high data quality and wide biological breadth, enabling systematic recognition of cell-selective elements. However, as genome-scale data from diverse cellular contexts have accumulated, systematic assessment of cell type- and state-selectivity has grown increasingly complex. It has become evident that large sets of DHSs that are widely distributed across the genome may nonetheless exhibit common (but complex) patterns of cell-selective actuation ^15^. However, the lack of a common coordinate system for DHSs has greatly hampered systematic identification and exploitation of these features.

Here we sought to expand a broad set of high-quality DHS maps, and to unify them into a common reference framework that both (i) incorporates precise genomic annotation that reflects observed biological variability in the pattern of DNA accessibility seen at individual elements, and (ii) captures complex cell-selective behaviors in a quantitative fashion. Both the unprecedented precision of genomic annotation and the richness with which cell-variable behaviors could be captured were directly enabled by joint analysis of high-quality data across hundreds of biological contexts, and thus unattainable from smaller scale data. We show that the integration of DNA accessibility maps across large numbers of cell and tissue states results in a remarkably coherent framework with wide utility for annotation of the human regulatory DNA
and gene landscapes; for defining how regulatory programs are encoded within the genome; and for clarifying links between genetic signals and genome-scale regulatory programs that
enable new insights into the organization and interpretation of non-coding disease- and trait-associated variation.

## A deep index of consensus human DHSs

To create deeply sampled reference maps of human regulatory DNA marked by DHSs, we performed DNase-seq^19^ on a wide range of human cell and tissue samples spanning all major human organ systems (**Fig. 1a**; **Methods**). Reference-grade data were created by rigorous quality screening for high signal-to-noise ratios and complex libraries (**Methods**), and were aggregated with prior high-quality data from the ENCODE^15^ and Roadmap Epigenomics ^32^ Projects. We conservatively selected 733 biosamples with high-quality data, representing 439 cell or tissue types and states (**Fig. 1a**; **Supplementary Table 1; Methods**). The vast majority of these data derived from primary ex vivo cell and tissue samples (72% of samples) and from primary cells in culture (11%), with the remainder (17%) derived from immortalized cell lines. Collectively these data represent a >6-fold expansion of cell types or states relative to the prior phase of ENCODE^15^.

**Figure 1.**
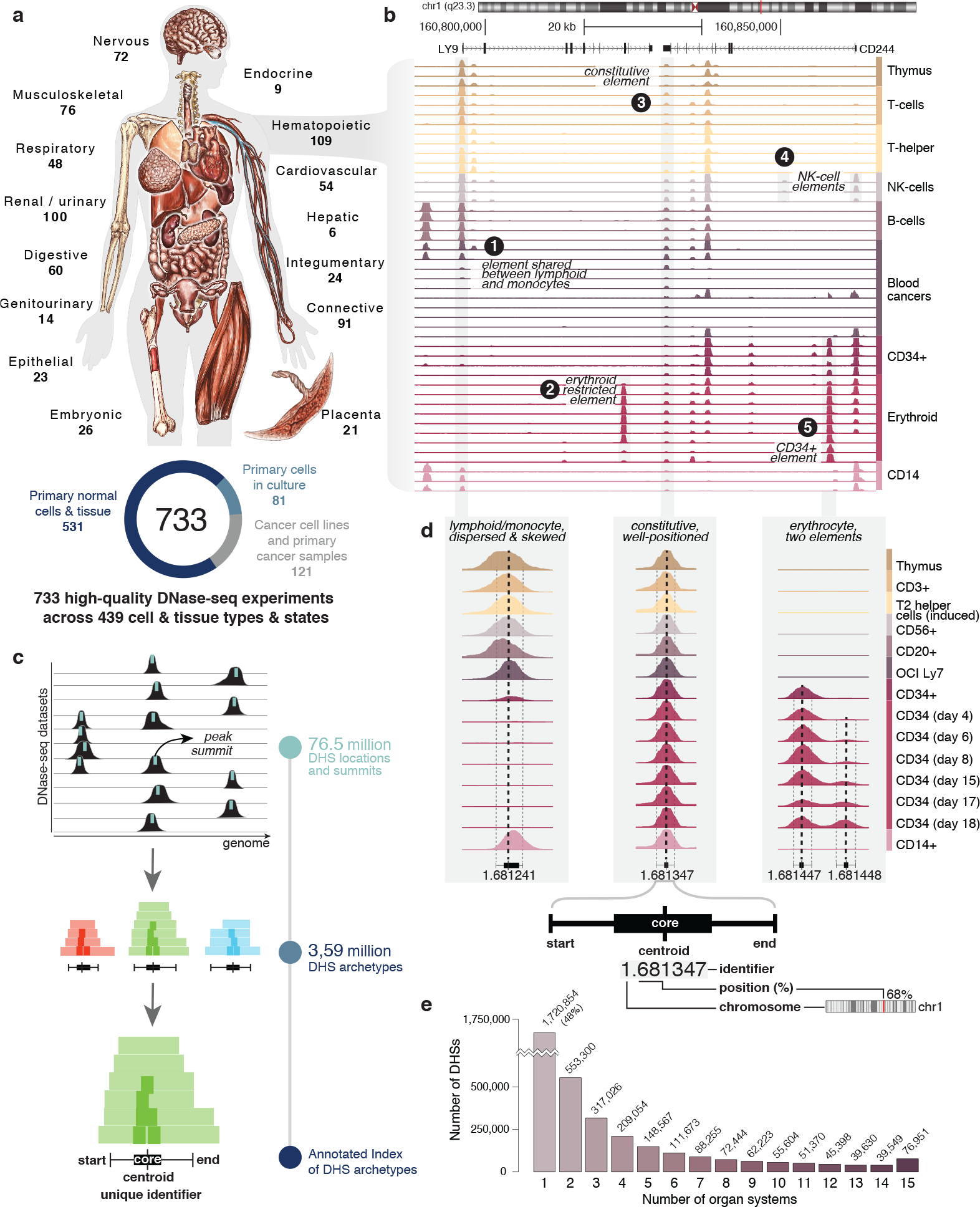
An index of DNase I Hypersensitive Sites (DHSs) in the human genome. **a.** DNA accessibility assayed across all main organ systems; number of datasets per system indicated. Out of 733 tota datasets, 531 are derived from primary cells and tissue. **b**. Example locus on chromosome 1, illustrating DNase I hypersensitivity across selected hematopoietic biosamples, as indicated on the right. A variety of cell type selective configurations are indicated. **c.** Outline of procedure to delineate a DHS Index; 76.5M per-dataset DHSs are aggregated across datasets to jointly delineate 3.59M consensus DHSs, with extensive annotation. **d.** Delineation of consensus DHSs shown for three example loci with varying cell type selectivity and positional stability. Positions of each per-dataset DHS are aggregated across datasets to determine consensus DHS coordinates, further annotated with a summit position, a core region reflecting the positional stability and a unique identifier. **e.** Sharing of DHSs across human organ systems, shown by the number of DHSs as a function of the number of organ systems in which they are observed.

### A common coordinate system for accessible regulatory DNA

Deeply sampled reference maps of DHSs reveal rich and varied patterns of DNase I hypersensitivity (Fig. 1b). We sought to create a precise and durable reference framework for genomic elements that encode DHSs by (i) comprehensively and stringently (FDR < 0.1%) delineating DHSs within each biosample using an improved algorithm (https://github.com/Altius/hotspot2); (ii) systematically integrating individual biosample maps to define archetypal DHS-encoding elements; and (iii) assigning to each archetypal DHS element a unique numerical identifier, thus establishing a common coordinate system for regulatory DNA marked by DHSs (Fig. 1c).

We identified an average of 104,433 DHSs per biosample (collectively detecting 76,549,656 DHSs across all 733 biosamples). To define archetypal DHS-encoding elements, we developed
a consensus approach outlined in **Fig. 1c** and **Extended Data Fig. 1a-b**. First, we aligned the summit coordinate (1bp) of each peak in DNase hypersensitivity signal across all biosamples to define the centroids of all spatially distinct DHSs. To resolve the boundaries of each element, we collated the local linear extent of DNase I hypersensitivity into a consensus range (**Methods**). We then combined centroids and boundaries into a single index of 3,591,898 distinct archetypal DHS-encoding sequence elements comprising, for each, (i) a consensus DHS summit; (ii) a ‘core’ region representing empirical confidence bounds on the centroid; and (iii) the consensus start and end coordinates of the archetypal DHS (**Fig. 1d**, **Extended Data Fig. 1c**). Because each DHS mapped within an individual biosample contributes to a single archetypal DHS in the consensus index, the provenance of each index DHS can be directly traced back to the DHSs of its contributing biosample(s). Finally, we assigned a unique identifier to each archetypal DHS within the index using a flexible and extensible numerical schema (**Fig. 1d**) that (i) conveys the genomic localization of each DHS; (ii) enables unlimited extension to newly-discovered DHSs; (iii) ensures compatibility with future reference genome builds and portability to personal genomes; and (iv) enables direct integration with DNase I footprints or other experimentally annotated components of a reference DHS (**Methods**). To create a robust framework for selecting more stringent genomic subsets of the complete index, we assigned confidence scores to all DHSs that reflect both the propensity for repeated observation in independent biosamples and the signal strength. These scores (**Extended Data Fig. 1d-e**) can be used to select DHS subsets at any desired level of stringency.

Index DHSs are broadly distributed across annotated genic and repetitive elements (**Extended Data Fig. 2a**). 54% of DHSs overlap repetitive elements, in line with observations that approximately half of human regulatory elements are derived from repetitive elements^33^, and covering all classes and sub(families) of repetitive elements (**Extended Data Fig. 2b**). Most DHSs localize distal to annotated TSSs (**Extended Data Fig. 2d**), 53% lie within introns, ~3% within non-coding exons and UTRs, and ~2% are dually encoded within protein-coding exons (**Extended Data Fig. 2c**).

Consensus indexing enabled a far more precise and compact annotation of the accessible regulatory DNA compartment of the human genome than previously attainable^15^. The ~3.6 million index DHSs have an average width of 204bp (median 196bp, interquartile range 151-240bp) and annotate 665.57 Mb or 21.55% of the human genome sequence. DHS ‘core’ regions within the index have an average width of 55bp (median 38bp) and annotate 197.74Mb or 6.4% of the genome. DHS summits precisely define the peak in evolutionarily conserved nucleotides within DHSs, and the corresponding trough in the density of human genetic variation (**Extended Data Fig. 2e**).

## Cellular patterning of DNA accessibility

DHSs are extensively shared across individual biosamples and, more generally, across groups of biosamples from different human organ systems (**Fig. 1e** and **Extended Data Fig. 2f**). Previously we described the existence of stereotyped cross-cell-type actuation patterns shared by tens to hundreds of widely distributed DHSs sharing the same biological function such as enhancer activity^15^. We observed a high degree of complexity in the patterning of index DHS actuation across the 733 cell types and states studied (**Fig. 2a**), with both biologically modular and less structured patterns (**Fig. 2b**). Across this very large sample set, DHSs are frequently selective for a single cell type or state, though a majority show complex actuation behavior across cell states (**Fig. 1e**, **Extended Data Fig. 2f**).

**Figure 2.**
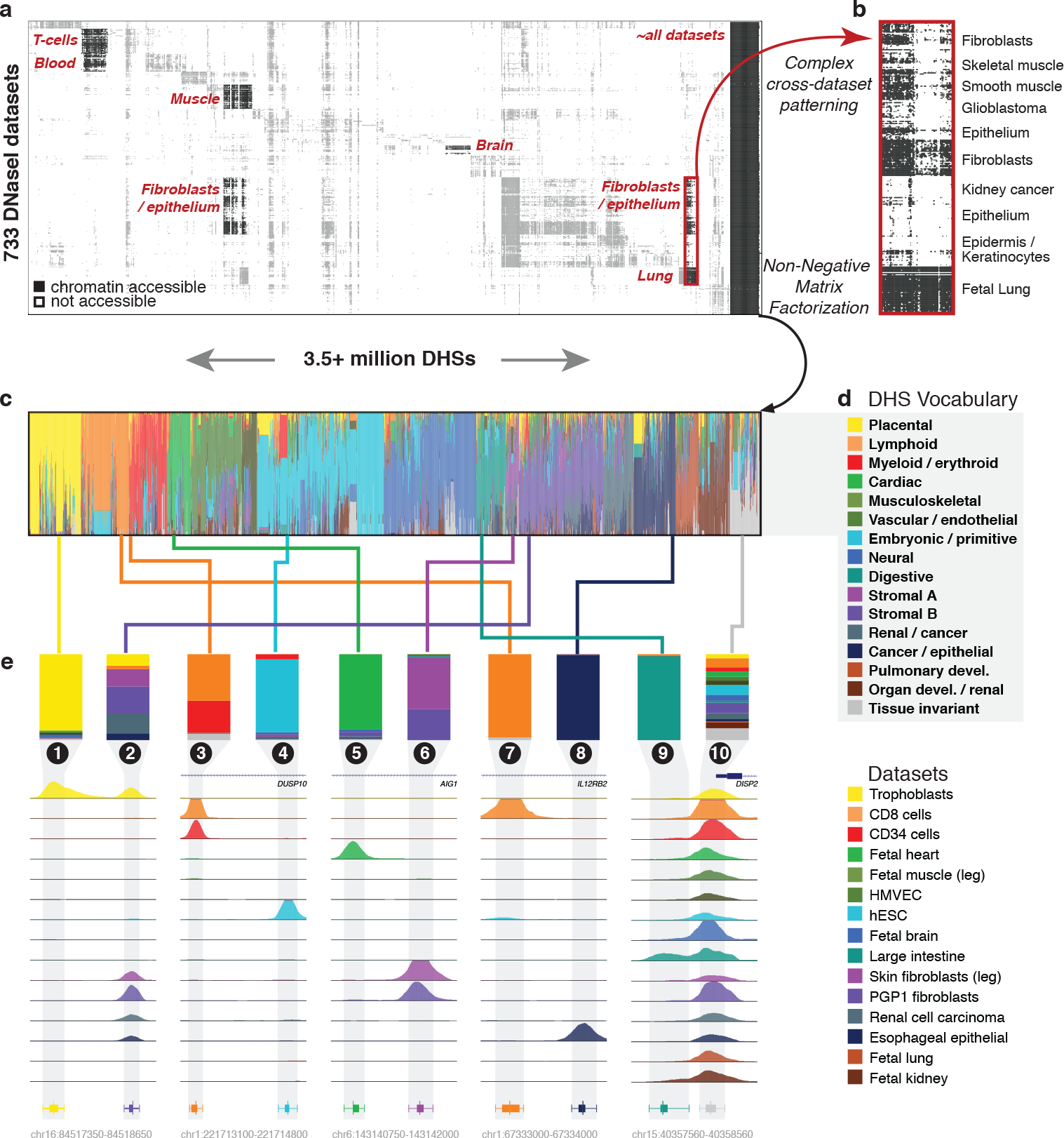
A simple vocabulary captures complex cross-dataset patterning of DHSs. **a.** DNA accessibility of 3.5+ million DHSs across 733 DNase-seq datasets, documented in a single DHS-by-dataset matrix. Indicated are various recurring accessibility patterns, including extensive sharing across cellular contexts. **b.** Example of a set of 1,000s of DHSs sharing similar accessibility patterns across cellular contexts, illustrating the modular behavior of DHSs. **c.** Decomposition of DHS patterns across 733 datasets into 16 components using Non-negative Matrix Factorization (NMF). The cellular patterning of individual DHSs is described using a mixture of components, indicated by distinct colors. **d.** Regulatory component labels, constituting a DHS vocabulary. **e.** Component mixtures for 10 exemplar DHSs, and corresponding DNase-seq data for representative cellular contexts. Shown are DHSs with various degrees of component-specificity, including a constitutive DHS shared across all indicated datasets and components. Single-component annotations representing dominant component loadings are shown at the bottom.

We next sought to develop a flexible approach for quantifying and annotating DHS actuation patterns. In principle, the cross-cell-state actuation pattern of any given index DHS can be summarized by a limited number of biological ‘components’ combined in a weighted fashion; the same components can be used (orthogonally) to summarize the DHS repertoire of an individual biosample. A key advantage of this approach is the possibility to capture complex behaviors while maintaining the potential for biological interpretability, since DHS-centric information can inform biosamples and *vice versa*.

### A compact vocabulary captures complex DHS regulatory patterns

To decompose the matrix of 3,591,898 DHSs × 733 biosamples we applied non-negative matrix factorization^34^ (NMF; **Extended Data Fig. 3a-c**), a technique initially employed in the field of computer vision to learn parts-based representations of objects, and semantic features of text^35^. We represented each DHS by a large enough number of components (*k*=16) to ensure accuracy – i.e., the degree to which the original matrix can be reconstructed from the components – while retaining potential for interpretability via assignment of components to established biological contexts such as known cell lineage relationships, or cell states known to be specified by specific regulatory factors (**Fig. 2c**, **Extended Data Fig. 3d-f, Methods**).

To connect components with established biological contexts, we identified (i) the biosamples most strongly associated with each component, and (ii) the distribution of TF recognition sequences within DHSs most strongly associated with that component. For all components, the top 15 contributing cell or tissue samples were remarkably coherent, enabling provisional assignment of a meaningful biological label to most components (**Extended Data Fig. 4a**). Beyond these top associations, more general analyses further support these labelings (**Extended Data Fig. 4b-d**, **Methods**).

We next analyzed the distribution and enrichment of TF recognition sequences within DHSs most strongly associated with each component (**Extended Data Fig. 5a**; **Methods**), revealing clear mappings between distinct sets of cell lineage- or state-specifying TFs and specific components (**Extended Data Fig. 5b**). Notably, these mappings are orthogonal to the biosample-to-component mappings described above. Finally, we combined biosample-to-component mappings and TF-to-component mappings to create a robust and biologically resonant regulatory ‘vocabulary’ that can be used to capture the actuation pattern of any DHS across cell types and states (**Fig. 2d**, **Box 1**).

### Biological annotation of DHSs using regulatory components

To date, over 99% of DHSs encoded by the human genome remain unannotated. Because the actuation pattern of each DHS across biosamples is captured by linear combinations of NMF components (**Fig. 2c**, **Extended Data Fig. 3c**), these combinations provide de facto annotations of the biological spectrum of every DHS (**Fig. 2e**). DHSs selective for a single cell type or state are annotated by a single majority component (**Fig. 2e**, columns 1,4,5,7,8,9); DHSs occurring in multiple cellular contexts are described by a combination of components (**Fig. 2e**, columns 2,3,6,10); and constitutive DHSs are annotated by a rich mix of all components (**Fig. 2e**, column 10), including a specific component that describes tissue-invariant behavior. In this schema, DHSs with similar cross biosample actuation patterns exhibit similar mixtures of components. For analytical practicality and visual compactness, the annotation of each DHS can be further summarized using its strongest single component (**Fig. 2e**, bottom); we use this summary vocabulary for the analyses described below.

## Regulatory annotation of human genes

The function of many genes is closely connected with their regulation across cells and tissues, and hence with the activity spectra of their cognate regulatory elements. We observed that DHSs with similar annotations were highly clustered along the genome (**Methods**; **Extended Data Fig. 6a-b**), particularly over gene bodies (**Extended Data Fig. 6c**). We thus reasoned that regulatory components of overlying DHSs could be utilized to annotate the likely functional compartments of their underlying genes. Quantifying the enrichment of congruently annotated DHSs around 56,832 GENCODE genes (protein-coding and non-coding) genome-wide revealed 20,658 genes (FDR < 5%) with significant clustering of DHSs belonging to the same component (**Fig. 3a-b**; **Supplementary Table 2**). This phenomenon was particularly striking for genes encoding tissue-regulatory factors such as GATA1 (myeloid/erythroid component DHSs), FOXP3 (lymphoid component), and HOXB9 (developmental component). A subset of genes showed enrichment of more than one component suggestive of different functions in different organ systems – for example CDX2 (embryonic/primitive and digestive components; **Fig. 3c**).

**Figure 3.**
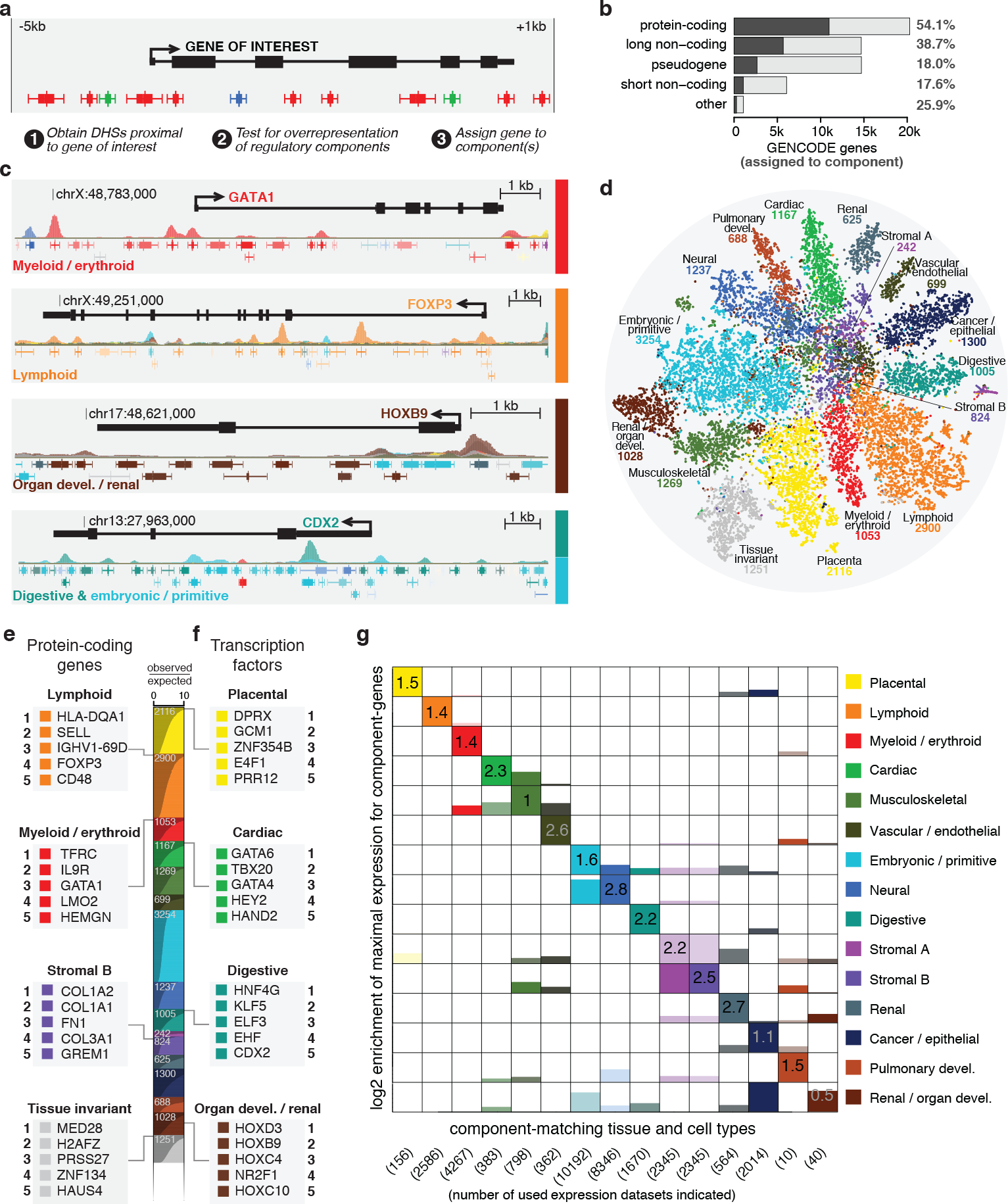
Regulatory annotation of human genes. **a.** Regulatory annotation of genes through the over-representation of regulatory components in a region defined by the body of a gene, extending a maximum of 5kb upstream and 1kb downstream, up until halfway through to another gene. **b**. Percentage of genes annotated with one or more regulatory components, split up by the major gene categories as recognized by GENCODE. **c.** Regulatory annotations of the protein-coding genes GATA1, FOXP3, HOXB9 and CDX2. **d.** 2D t-SNE projection of regulatory annotation patterns across genes, colored by their most strongly associated component. The number of genes associated with each regulatory component is indicated. **e-f.** Summarized view of the number of genes associated with each regulatory component. Call-outs show the top 5 results for protein-coding genes (**e**; myeloid/erythroid, lymphoid, stromal and tissue-invariant components) and the subset of transcription factor genes (**f**; placenta, cardiac, digestive or organ development / renal components). **g.** Correspondence between gene regulatory annotation and cell type of maximal RNA expression shown using relative transcriptional activity of genes in a panel of component-matched tissue and cell types. Values shown are log2 observed/expected ratios.

Of 20,291 GENCODE protein coding genes, more than half (54.1%) could be assigned a regulatory component based on their overlying DHSs (**Fig. 3b**). To determine whether these assignments are concordant with their annotated function, we assessed (i) whether the most confidently annotated genes reflect their known function and (ii) whether genes annotated with a particular component are maximally expressed in cell types matching those components. The top genes annotated by the lymphoid component are all involved in immune response and disease (**Fig. 3e**, **Extended Data Fig. 7a**). Similar relationships were observed for other categories of genes including those annotated by the myeloid/erythroid component (erythropoiesis or hematopoietic stem cell genes, **Fig. 3e**, **Extended Data Fig. 7b**), a stromal component (collagen genes and fibronectin, **Fig. 3e**, **Extended Data Fig. 7c**), and the tissue-invariant component (housekeeping genes, **Fig. 3e**, **Extended Data Fig. 7d**). This phenomenon was particularly striking for TF-encoding genes ^36^ such as lineage specifying master regulators for cardiac development (cardiac component, **Fig. 3f**, **Extended Data Fig. 8b**) or development of other organ systems (**Extended Data Fig. 8**).

To explore the concordance between regulatory vocabulary annotations and gene expression across cell types and states, we interrogated a compendium of over 100,000 uniformly processed RNA-seq data sets^37^. After matching regulatory components with tissue-relevant expression data sets (Methods), we observed strong correspondence between the vocabulary-based annotation of genes and the cell or tissue types in which they were maximally expressed (Fig. 3g).

### Annotation of genes with unknown functions

Despite intensive study, the function of many human genes remains obscure, particularly those with lowly or highly cell-selective expression patterns – for example, zinc-finger (ZNF) TFs^36,38^ or long non-coding RNA genes^39^. Nearly half of ZNF TFs (43.7%) could be annotated with a single regulatory component (**Extended Data Fig. 9a**) thus indicating their likely target biological sphere of activity. 38.7% of long non-coding genes evinced clear mappings to regulatory components (**Extended Data Fig. 9b**), as did 18% of pseudo-genes^40^ (**Extended Data Fig. 9c**), possibly reflective of remnants of regulatory states prior to ancient gene duplications. Beyond genes, we reasoned that entire pathways could be annotated using the overlying DHS landscapes of their constituent genes (**Extended Data Fig. 10a**) – for instance, the KEGG pathway ‘Allograft rejection’ is strongly enriched for the lymphoid component (**Extended Data Fig. 10b**), consistent with the principle that genes involved in similar biological processes share similar patterns of regulatory element actuation.

## Annotation of trait-associated genetic variation

We next asked whether DHS annotations could expand insights into the role(s) of genetic variation harbored within regulatory DNA, and provide a more meaningful framework for interpreting the pathophysiological basis of disease and trait associations. A rank-based analysis of disease/trait versus regulatory component associations (explicitly controlling for large scale Linkage Disequilibrium (LD) structure; Methods) revealed increasingly strong component-specific enrichments of association signals across diverse traits (**Fig. 4a**, **Extended Data Fig. 11a,b**). Notably, the observed enrichments could not typically be obtained by considering only DHSs from biosamples most closely related to the relevant regulatory component (e.g., lymphoid cell biosamples vs. lymphoid component; **Fig. 4a**, **Extended Data Figs. 4a, 11c**).

**Figure 4.**
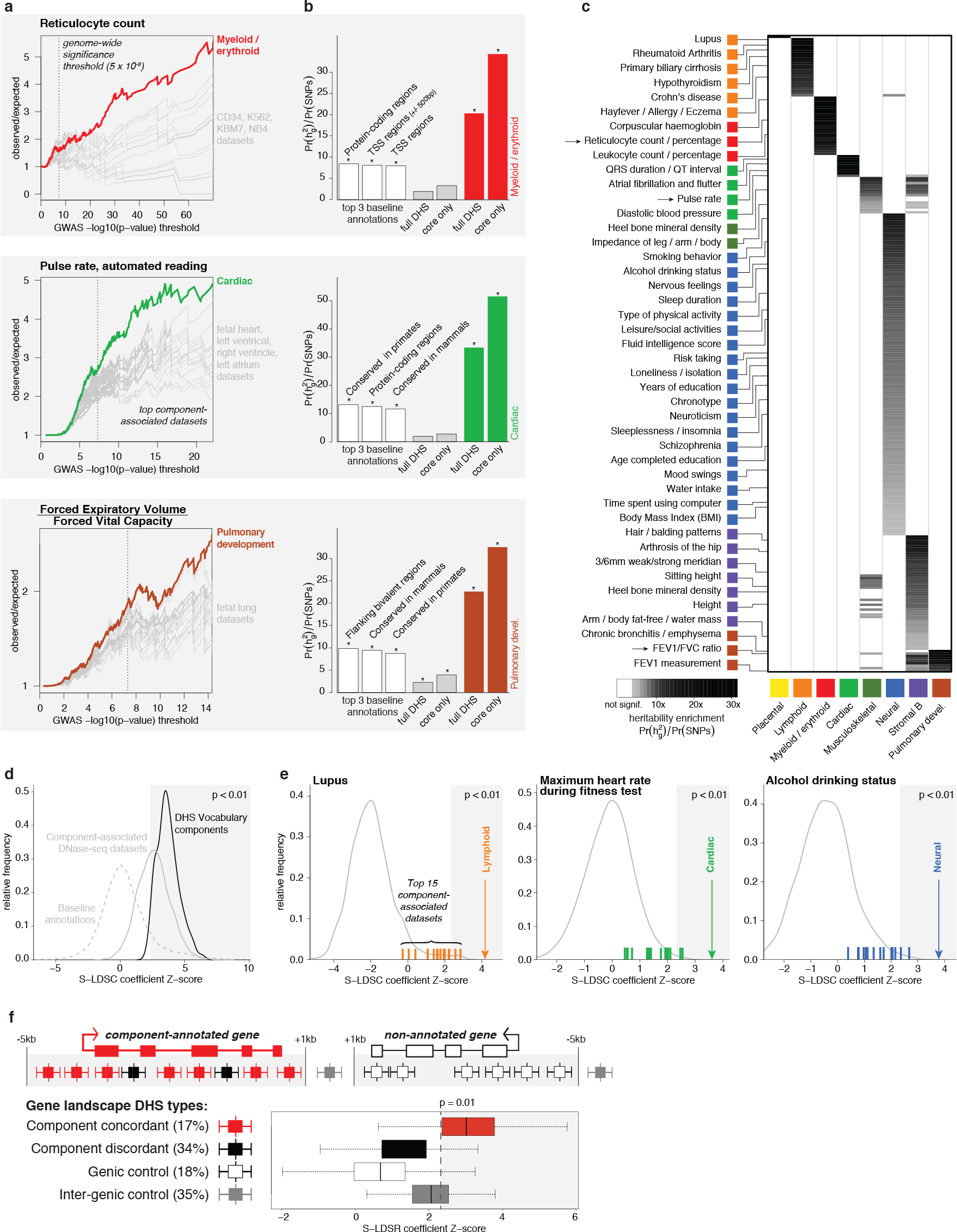
Component-wise view of disease-associated genetic variation. **a.** Per-component association of DHSs with selected GWAS traits, shown as enrichment ratios as a function of increasingly stringent subsets of variants. The canonical genome-wide significance threshold (5 × 10-8) is indicated. Enrichments for the top 15 biosamples associated with each regulatory component are shown in grey. **b.** Associations between GWAS variants and regulatory components identified through stratified LD-score regression (S-LDSC), for the traits shown in **a.** Heritability enrichment levels are indicated for the top 3 most enriched baseline annotations (white), the full DHS index (grey) and trait-relevant regulatory components (colored bars). Statistically significant enrichments are indicated (*; FDR < 0.01). **c.** Regulatory component (x-axis) heritability enrichment results across 261 GWAS traits (y-axis). Grayscale intensity indicates heritability enrichment levels, shown for statistically significant associations at FDR < 1%. A sampling of labels of enriched traits is shown for each component, as space permits. Traits used in **a** and **b** are indicated with arrows. **d.** Distribution of S-LDSC coefficient Z-scores across 261 GWAS traits, shown for all baseline annotations (grey dashed line), top 15 regulatory component-associated biosamples (grey solid line) and regulatory components (black line). **e.** S-LDSC coefficient Z-scores for selected traits, shown for all individual biosamples (grey lines), top 15 component-associated biosamples (colored ticks) and regulatory components (colored arrows). **f.** S-LDSC Z-scores stratified according to gene landscape DHS types, indicating stronger heritability contributions for component concordant DHSs. For **d-f**, Grey areas indicate S-LDSC Z-scores corresponding to p < 0.01.

Quantifying the extent to which DHS annotations captured SNP-based trait heritability (h_g_^2^, **Fig. 4b**, white bars) revealed a strong increase in heritability enrichment for trait-relevant regulatory components relative to index DHSs (**Fig. 4b**, colored bars vs. grey bars, respectively). Heritability was markedly enriched specifically within DHS ‘core’ regions, providing orthogonal evidence supporting the delineation and importance of these subregions (**Fig. 4b**).

To generalize these observations, we compiled >1,300 traits with SNP-based heritability of at least 1% from the UK Biobank project^41^ and from curated published data^42^. Of these, 261 diseases and traits showed highly significant component-specific enrichment in heritability, particularly for pathophysiologically relevant regulatory components (**Fig. 4c**, **Extended Data Fig. 12a**, FDR < 1%). Restricting DHS delineations to ‘core’ regions again yielded significantly greater enrichment compared to full DHSs (**Extended Data Fig. 12b-c**). To remove potentially confounding contributions from multiple genomic annotations overlapping the same SNP, we quantified the statistical significance of regulatory component heritability contributions while controlling for the contribution of all other annotations (**Methods**). For virtually all reported traits, regulatory component annotations significantly (p < 0.01) captured SNP-based trait heritability (**Fig. 4d**, black line).

To quantify the concentration of trait-associated genetic signal in DHSs annotated by specific regulatory components relative to the full repertoiretype-specific heritabil of DHSs in disease/trait-relevant cell types we performed cell ity analyses^43^ (**Methods**). Component-annotated DHSs offer a significant improvement in capturing trait heritability compared to individual datasets (p < 2.2 × 10^−16^; **Fig. 4d** grey solid line). Strikingly, we find that at the level of individual traits, in 68 out of 261 (26%) cases regulatory component annotations better capture trait heritability than individual DNase-seq datasets (**Fig. 4e**).

### Concentration of trait-associated variation in component-annotated gene body DHSs

The clustering of concordantly regulated DHSs within gene bodies (**Fig. 3**) prompted us to speculate that such DHSs were more likely to contain relevant genetic signals. To test this, we quantified the significance of regulatory component heritability separately for concordant (17% of DHSs) and discordant (34%) DHS annotations (**Fig. 4f**). Although in the minority, the former uniquely and strongly contributed to SNP-based trait heritability relative to DHSs occurring in the same genes but discordant with the gene’s component labeling. DHSs proximal to genes not labelled by any regulatory component provide the weakest heritability contributions, while intergenic DHSs contribute modestly (**Fig. 4f**, **Extended Data Fig. 12d**).

Taken together, our results show that partitioning the trait heritability encoded in DHSs based on regulatory components provides a novel and powerful approach for prioritization of genetic signals.

## Discussion

Here we have presented by far the most comprehensive and precise map of human DNase I hypersensitive sites, the best described generic markers of regulatory DNA. The remarkable positional stability of DHSs across cell types and states has enabled delineation and indexing of archetypal DHS-encoding sequence elements within the human genome. We have also provided the first universal annotation that captures, for every archetypal DHS, its complex actuation pattern across diverse cell types and states. Together these features create a powerful new framework for analyses at the intersection of gene regulation and the genetics of human diseases and quantitative traits. Archetypal DHS identifiers are robust to genome builds, transferable to personal genomes and emerging graph-based genome analysis^44,45^, and enable facile incorporation of functional properties such as cell-selectivity, or finer structural annotations such as DNase I footprints. Common reference coordinates will further greatly facilitate comparisons between large experimental data sets, and between human and mouse DHSs.

Cell type-selectivity is a cardinal property of DHSs yet has been difficulty to analyze systematically or to compare between organisms and individuals due to lack of a common positional reference framework. Regulatory components capture complex cell context behaviors and thus greatly expand the analytical horizon beyond cell type-agnostic annotations such as chromatin states^30,46^. Because each regulatory component can be mapped to a set of contributing TFs and resolved to specific archetypal DHSs, they provide annotations of unprecedented richness that can be readily leveraged for mechanistic insights into regulatory pathways and networks.

By combining DHS indexing and regulatory components, it is now possible to triangulate the genetics-gene regulation interface on three axes: (i) a genomic position axis, which has been finely resolved to consensus DHS summits; (ii) a cell/tissue biological axis captured in regulatory components; and (iii) a gene context axis reflecting the coherent co-localization of similarly regulated DHSs over gene bodies.

On the genomic position axis, the concentration of trait heritability within highly compact ~50bp DHS cores – even relative to the immediately adjacent DHS ‘arms’ – is particularly striking given that the overall density of human sequence variation nadirs at DHS summits (**Extended Data Fig. 2e**). On the cell/tissue biological axis, regulatory components now provide a systematic and principled approach for combining the complex biological landscapes of individual cell types, resulting in a more powerful and comprehensive framework for analyzing the intersection of cell/tissue selectivity and trait heritability than could be achieved with individual biosamples or simple combinations thereof. Finally, on the gene axis, the concentration of GWAS variants within DHSs sharing a gene’s dominant component reinforces the biological coherence of these elements and provides a new pathway for separating variants implicating a given gene from those within its body or flanking regions that are connected with nearby or distant genes.

On a broader level, the DHS index and component framework we report represents a transition from an exploratory era focused on discovery of novel elements, to a map-centric framework with a focus on detection of an annotated element within a particular biological context. This new framework should prove particularly valuable for anchoring single cell studies, which are presently at least 1000-fold too sparse for robust delineation of DHSs within individual cells. The index/component framework is highly information rich, and provides an inter-operable reference for annotating functional connections between and among DHSs and genes, and for functional categorization of elements – two intertwined challenges that represent the next frontier in human genome annotation.

#### Box 1: DHS vocabulary component labeling

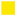**Placenta and trophoblast biosamples** – Strong enrichment for binding sites of GCM1, selectively expressed in trophoblasts and placenta in general^47^ and associated with pre-eclampsia in pregnant women^48^.

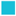**Primitive/embryonic** – Embryonic stem cell and related biosamples. Most enriched motif is that for POU5F1/OCT4, reflecting its key role in embryonic development and pluripotency.

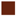**Organ development/renal** – Largely captures fetal kidney biosamples, with strong enrichment for development-related Homeobox protein (HOX) factors, as well as PAX2, associated with early kidney development^49^.

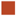**Pulmonary development** – Consists of fetal lung biosamples. Enriched for CEBPB and FOXC2 motifs, the latter of which is implicated in lung development and maturation^50^.

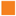**Lymphoid** – T-cells and other immune-related cellular conditions. Its most enriched motifs are Interferon-Regulatory Factors (IRF4, IRF1, IRF5, etcetera), in line with their critical role in the (adaptive) immune system^51^.

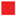**Myeloid/erythroid** – CD34+ cells and shows strong motif enrichments for not only ETS/SPI1, but also for GATA1 - by itself, as well together with TAL1.

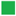**Cardiac** – Associated with heart-related biosamples. Strongly enriched for motifs of the Myocyte enhancer factor-2 (MEF2) transcription factor, a core cardiac transcription factor^52^.

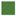**Musculoskeletal** – Associated with muscle(-related) and bone biosamples. Enriched for Musculin (MSC) motifs, with known key roles in regulating myogenesis.

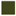**Vascular/endothelial** – Consists mostly of HMVEC cells and is enriched for motifs of ERG, a member of the erythroblast transformation-specific (ETS) family of transcription factors, required for platelet adhesion to the subendothelium, inducing vascular cell remodeling.

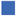**Neural** – Brain and other nervous system biosamples. General enrichment for AT-rich homeobox motifs, as well as many highly specific NEUROD2 motifs.

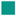**Digestive** – Associated with intestine, liver and bowel mucosa biosamples. Strong enrichment for motifs of hepatocyte nuclear factor 4 (HNF4), critical for liver development.

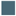**Renal/cancer** – Mostly adult kidney biosamples, including renal cancer. Enriched for HNF1B, the HNF1A paralog involved in kidney function.

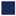**Cancer/epithelial** – Associates with various cancer types, as well as general epithelia. The top-scoring motif is for p53 (cancer-related) and p63, which has been proposed to play a dual role^53^: initiating epithelial stratification during development and maintaining proliferative potential of basal keratinocytes in mature epidermis.

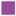**Stromal A** – Captures fibroblast biosamples. Enriched for motifs of Jun dimerization protein 2 (JDP2), as well as other components of the AP-1 transcription factor (JUN, FOS).

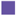**Stromal B** – Captures similar Biology as the Stromal A component.

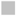**Tissue invariant** – Lacks strong association with specific cell and tissue types, but shows enrichment for a diverse set of housekeeping factor motifs, such as CTCF, ETS and NRF.

