## Supplemental Figures for "Index and biological spectrum of accessible DNA elements in the human genome"

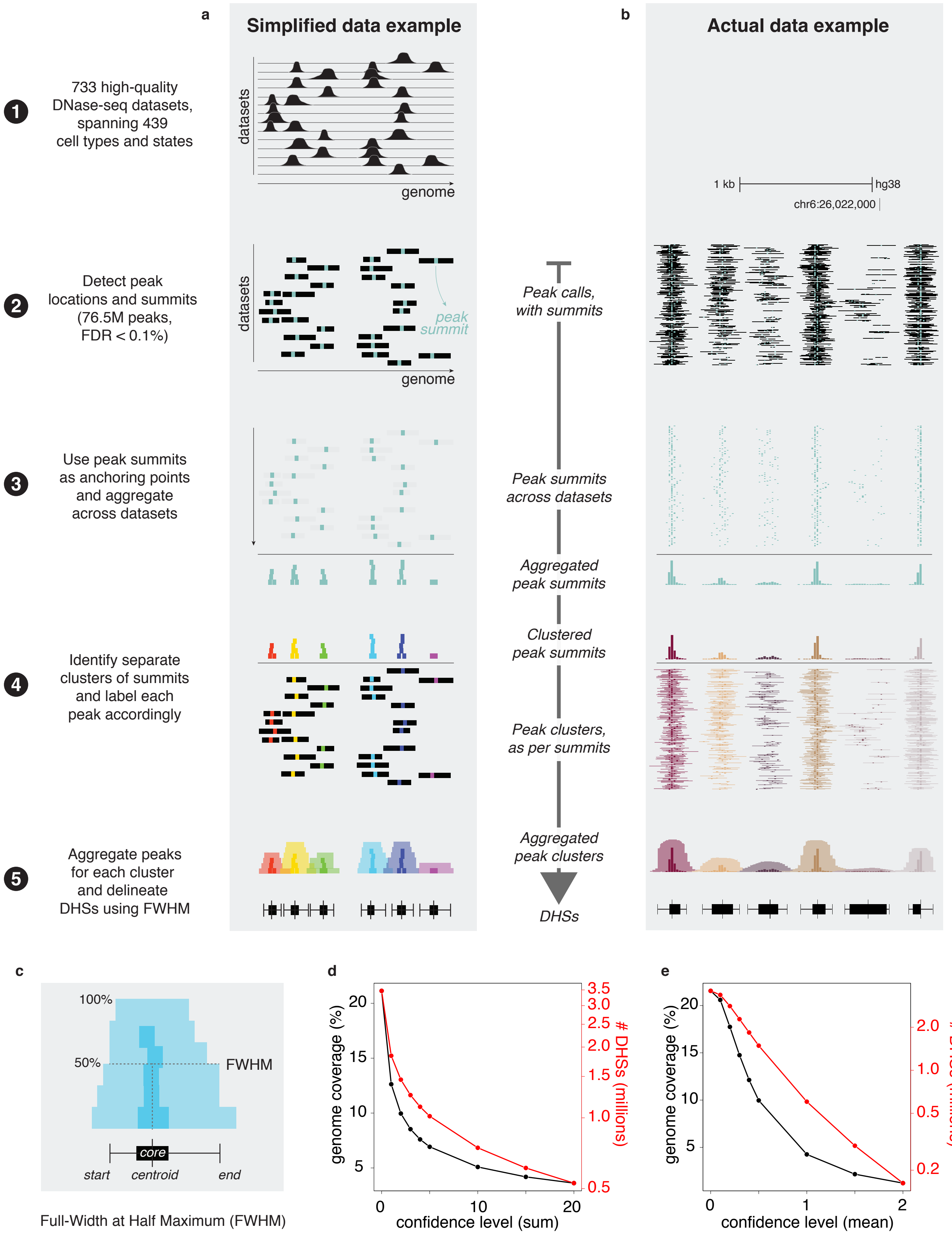

a
Simplified data example
b
Actual data example
1
733 high-quality DNase-seq datasets, spanning 439 cell types and states
datasets
genome
1 kb
hg38
chr6:26,022,000
2
Detect peak locations and summits (76.5M peaks, FDR < 0.1%)
peak summit
Peak calls, with summits
3
Use peak summits as anchoring points and aggregate across datasets
Peak summits across datasets
Aggregated peak summits
Clustered peak summits
4
Identify separate clusters of summits and label each peak accordingly
Peak clusters, as per summits
5
Aggregate peaks for each cluster and delineate DHSs using FWHM
Aggregated peak clusters
DHSs
c
100%
50%
FWHM
core
start
centroid
end
Full-Width at Half Maximum (FWHM)
d
genome coverage (%)
### DHSs (millions)
confidence level (sum)
e
confidence level (mean)


ED Figure 1

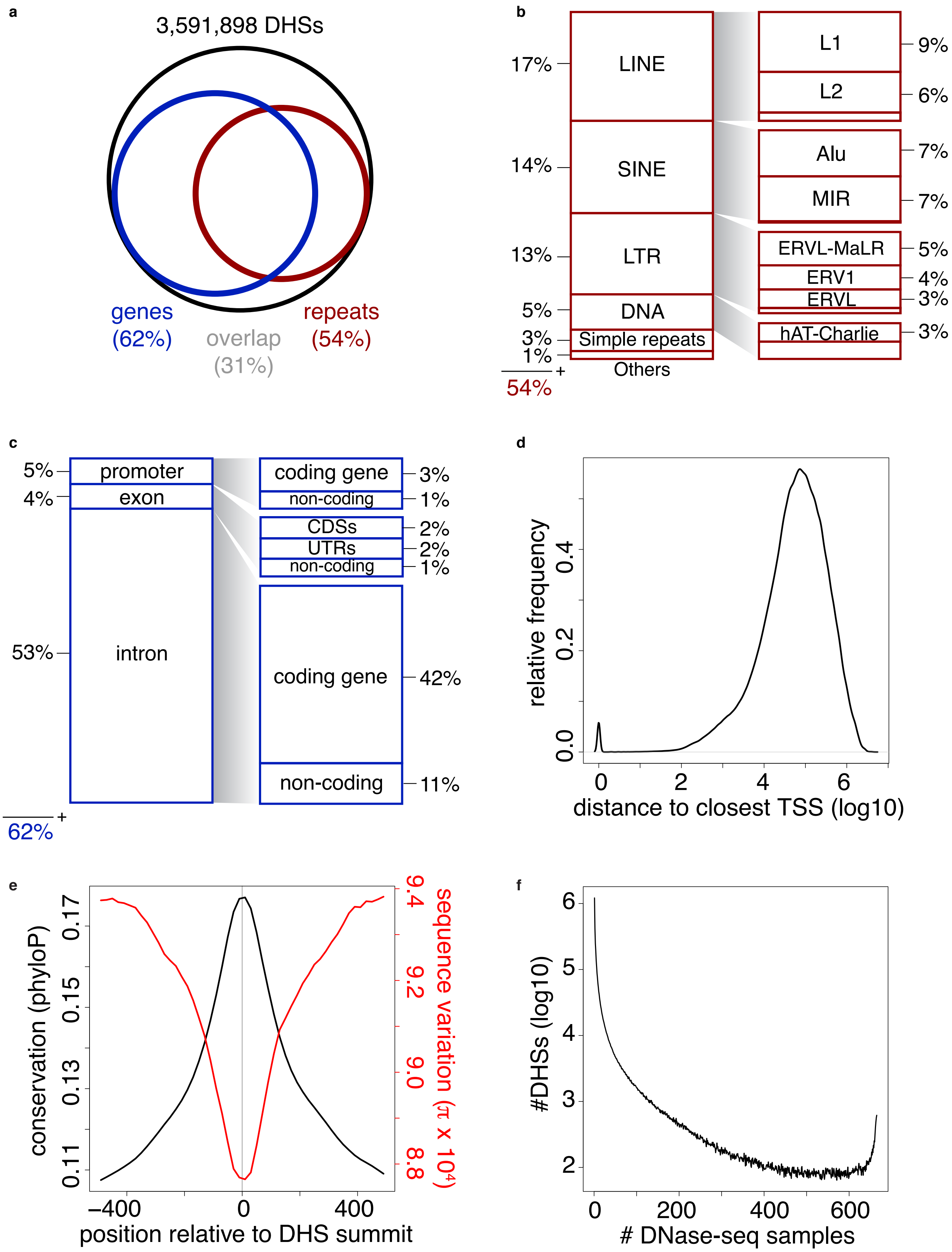
a
3,591,898 DHSs
genes (62%)
overlap (31%)
repeats (54%)
b
17%
LINE
14%
SINE
13%
LTR
5%
DNA
3%
Simple repeats
1%
Others
54%
L1
9%
L2
6%
Alu
7%
MIR
7%
ERVL-MaLR
5%
ERV1
4%
ERVL
3%
hAT-Charlie
3%
c
5%
promoter
4%
exon
53%
intron
62%
coding gene
3%
non-coding
1%
CDSs
2%
UTRs
2%
non-coding
1%
coding gene
42%
non-coding
11%
d
relative frequency
distance to closest TSS (log10)
e
conservation (phyloP)
sequence variation (π x 10⁴)
position relative to DHS summit
f
#DHSs (log10)
### DNase-seq samples

ED Figure 2

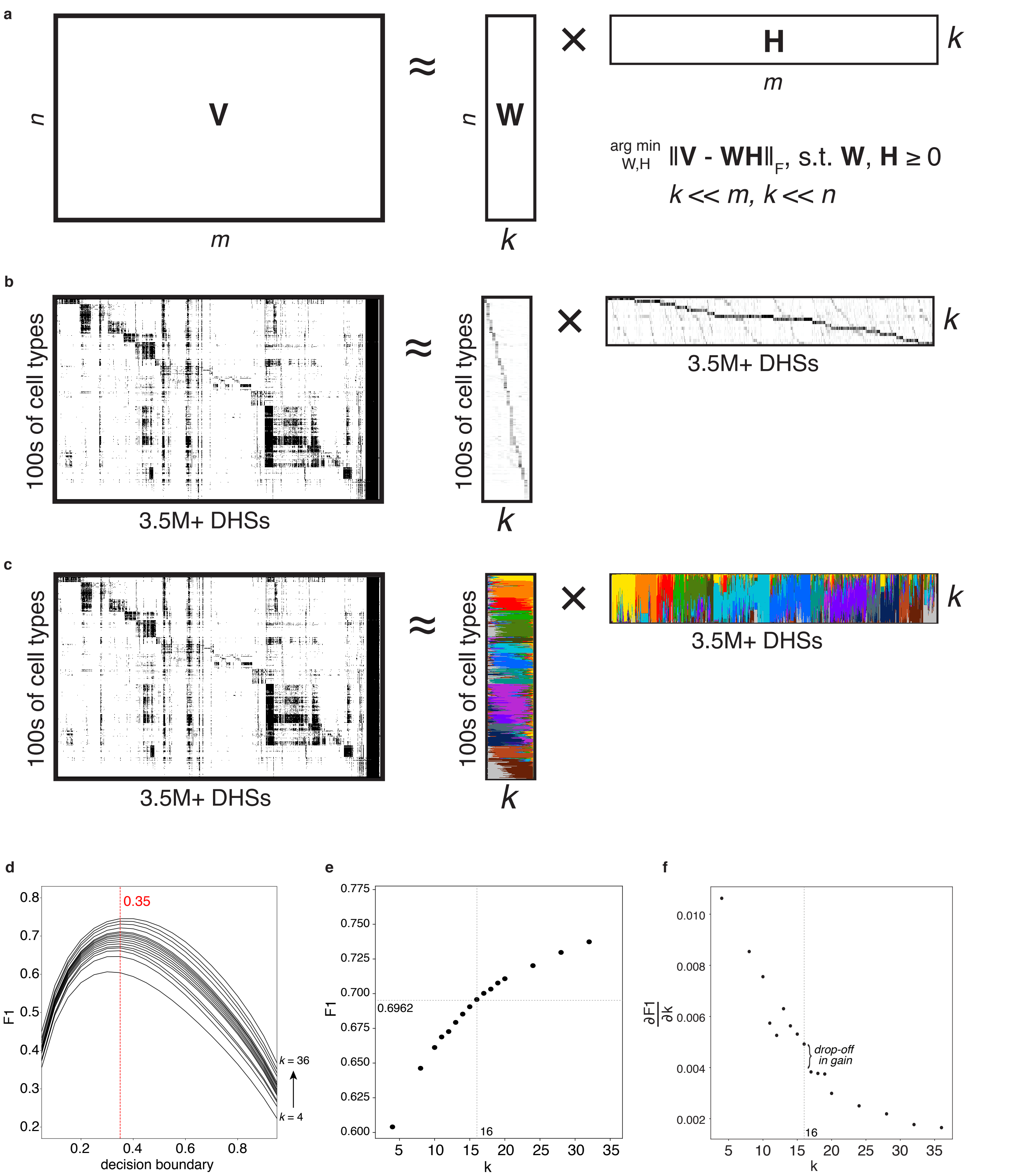
a
V
n
m
≈
W
n
k
×
H
k
m
arg min W,H ‖V - WH‖F, s.t. W, H ≥ 0
k << m, k << n
b
100s of cell types
3.5M+ DHSs
k
c
d
F1
decision boundary
0.35
k = 36
k = 4
e
0.6962
16
k
f
∂F1/∂k
drop-off in gain

ED Figure 3

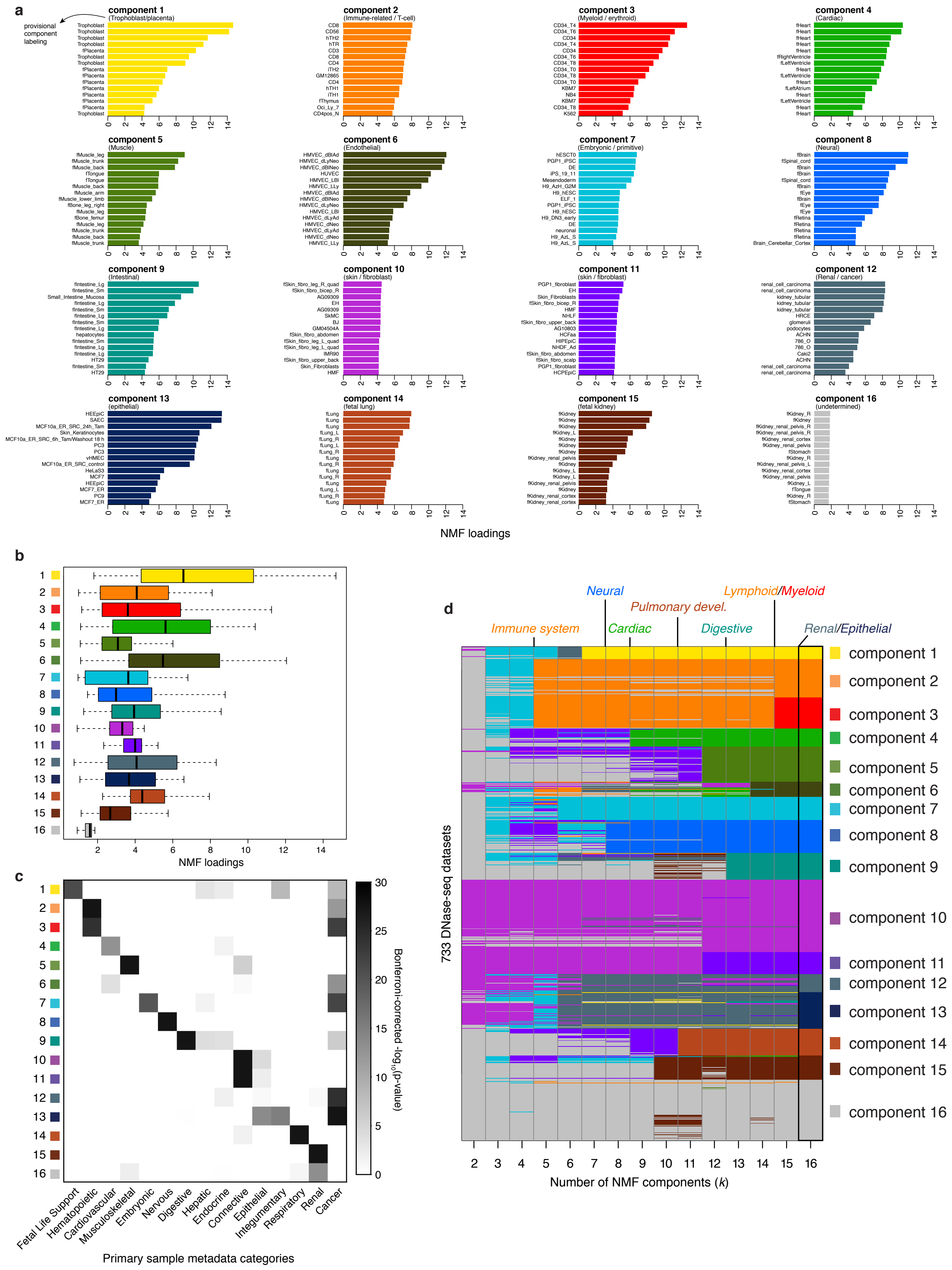
a
provisional component labeling
component 1
(Trophoblast/placenta)
component 2
(Immune-related / T-cell)
component 3
(Myeloid / erythroid)
component 4
(Cardiac)
component 5
(Muscle)
component 6
(Endothelial)
component 7
(Embryonic / primitive)
component 8
(Neural)
component 9
(Intestinal)
component 10
(skin / fibroblast)
component 11
(skin / fibroblast)
component 12
(Renal / cancer)
component 13
(epithelial)
component 14
(fetal lung)
component 15
(fetal kidney)
component 16
(undetermined)
NMF loadings
b
NMF loadings
c
Bonferroni-corrected -log10(p-value)
Fetal Life Support
Hematopoietic
Cardiovascular
Musculoskeletal
Embryonic
Nervous
Digestive
Hepatic
Endocrine
Connective
Epithelial
Integumentary
Respiratory
Renal
Cancer
Primary sample metadata categories
d
Neural
Lymphoid/Myeloid
Pulmonary devel.
Immune system
Cardiac
Digestive
Renal/Epithelial
733 DNase-seq datasets
component 1
component 2
component 3
component 4
component 5
component 6
component 7
component 8
component 9
component 10
component 11
component 12
component 13
component 14
component 15
component 16
Number of NMF components (k)

ED Figure 4

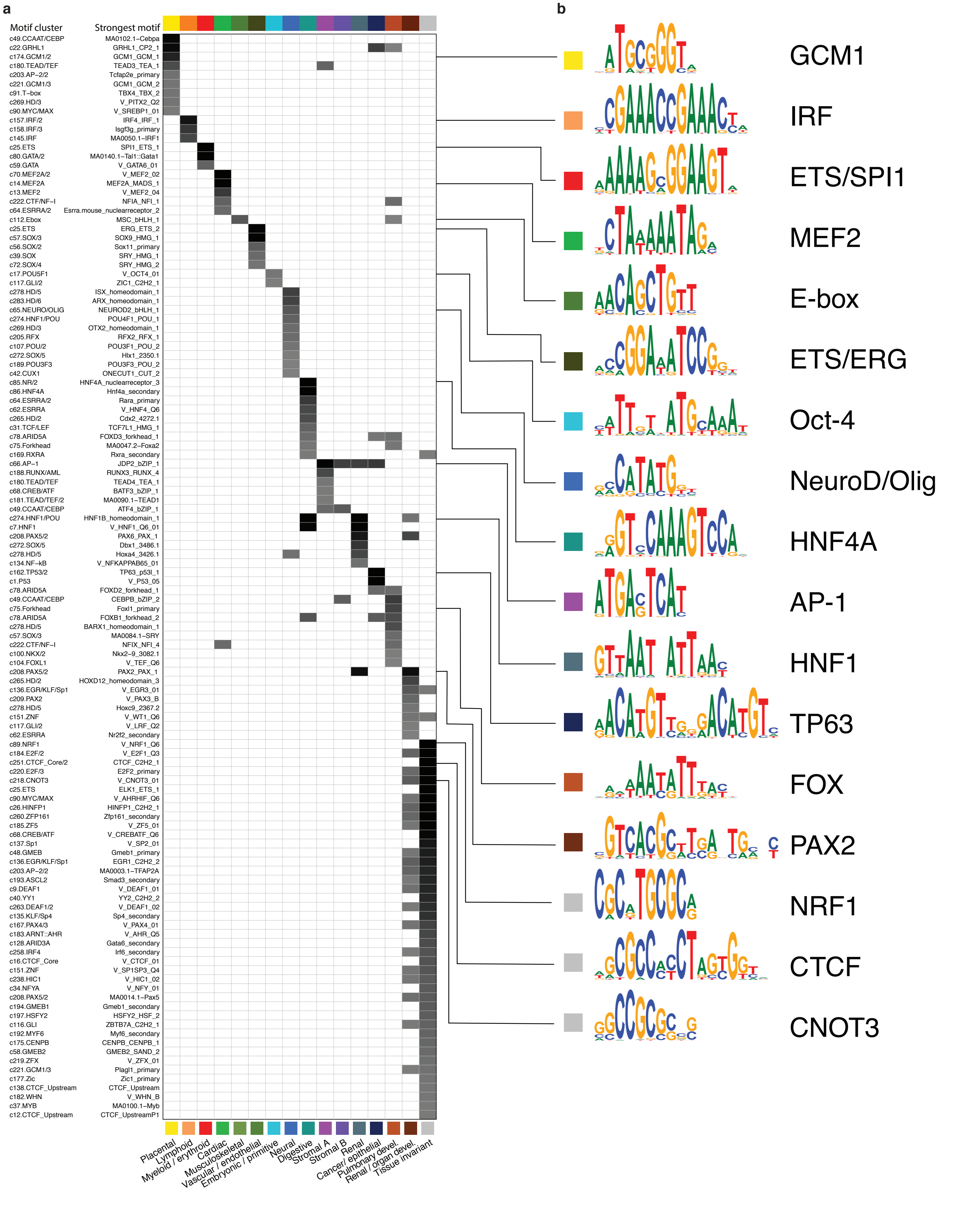
a
Motif cluster
Strongest motif
c49.CCAAT/CEBP MA0102.1–Cebpa
c22.GRHL1 GRHL1_CP2_1
c174.GCM1/2 GCM1_GCM_1
c180.TEAD/TEF TEAD3_TEA_1
c203.AP–2/2 Tcfap2e_primary
c221.GCM1/3 GCM1_GCM_2
c91.T–box TBX4_TBX_2
c269.HD/3 V_PITX2_Q2
c90.MYC/MAX V_SREBP1_01
c157.IRF/2 IRF4_IRF_1
c158.IRF/3 Isgf3g_primary
c145.IRF MA0050.1–IRF1
c25.ETS SPI1_ETS_1
c80.GATA/2 MA0140.1–Tal1::Gata1
c59.GATA V_GATA6_01
c70.MEF2A/2 V_MEF2_02
c14.MEF2A MEF2A_MADS_1
c13.MEF2 V_MEF2_04
c222.CTF/NF–I NFIA_NFI_1
c64.ESRRA/2 Esrra.mouse_nuclearreceptor_2
c112.Ebox MSC_bHLH_1
c25.ETS ERG_ETS_2
c57.SOX/3 SOX9_HMG_1
c56.SOX/2 Sox11_primary
c39.SOX SRY_HMG_1
c72.SOX/4 SRY_HMG_2
c17.POU5F1 V_OCT4_01
c117.GLI/2 ZIC1_C2H2_1
c278.HD/5 ISX_homeodomain_1
c283.HD/6 ARX_homeodomain_1
c65.NEURO/OLIG NEUROD2_bHLH_1
c274.HNF1/POU POU4F1_POU_1
c269.HD/3 OTX2_homeodomain_1
c205.RFX RFX2_RFX_1
c107.POU/2 POU3F1_POU_2
c272.SOX/5 Hlx1_2350.1
c189.POU3F3 POU3F3_POU_2
c42.CUX1 ONECUT1_CUT_2
c85.NR/2 HNF4A_nuclearreceptor_3
c86.HNF4A Hnf4a_secondary
c64.ESRRA/2 Rara_primary
c62.ESRRA V_HNF4_Q6
c265.HD/2 Cdx2_4272.1
c31.TCF/LEF TCF7L1_HMG_1
c78.ARID5A FOXD3_forkhead_1
c75.Forkhead MA0047.2–Foxa2
c169.RXRA Rxra_secondary
c66.AP–1 JDP2_bZIP_1
c188.RUNX/AML RUNX3_RUNX_4
c180.TEAD/TEF TEAD4_TEA_1
c68.CREB/ATF BATF3_bZIP_1
c181.TEAD/TEF/2 MA0090.1–TEAD1
c49.CCAAT/CEBP ATF4_bZIP_1
c274.HNF1/POU HNF1B_homeodomain_1
c7.HNF1 V_HNF1_Q6_01
c208.PAX5/2 PAX6_PAX_1
c272.SOX/5 Dbx1_3486.1
c278.HD/5 Hoxa4_3426.1
c134.NF–kB V_NFKAPPAB65_01
c162.TP53/2 TP63_p53l_1
c1.P53 V_P53_05
c78.ARID5A FOXD2_forkhead_1
c49.CCAAT/CEBP CEBPB_bZIP_2
c75.Forkhead Foxl1_primary
c78.ARID5A FOXB1_forkhead_2
c278.HD/5 BARX1_homeodomain_1
c57.SOX/3 MA0084.1–SRY
c222.CTF/NF–I NFIX_NFI_4
c100.NKX/2 Nkx2–9_3082.1
c104.FOXL1 V_TEF_Q6
c208.PAX5/2 PAX2_PAX_1
c265.HD/2 HOXD12_homeodomain_3
c136.EGR/KLF/Sp1 V_EGR3_01
c209.PAX2 V_PAX3_B
c278.HD/5 Hoxc9_2367.2
c151.ZNF V_WT1_Q6
c117.GLI/2 V_LRF_Q2
c62.ESRRA Nr2f2_secondary
c89.NRF1 V_NRF1_Q6
c184.E2F/2 V_E2F1_Q3
c251.CTCF_Core/2 CTCF_C2H2_1
c220.E2F/3 E2F2_primary
c218.CNOT3 V_CNOT3_01
c25.ETS ELK1_ETS_1
c90.MYC/MAX V_AHRHIF_Q6
c26.HINFP1 HINFP1_C2H2_1
c260.ZFP161 Zfp161_secondary
c185.ZF5 V_ZF5_01
c68.CREB/ATF V_CREBATF_Q6
c137.Sp1 V_SP2_01
c48.GMEB Gmeb1_primary
c136.EGR/KLF/Sp1 EGR1_C2H2_2
c203.AP–2/2 MA0003.1–TFAP2A
c193.ASCL2 Smad3_secondary
c9.DEAF1 V_DEAF1_01
c40.YY1 YY2_C2H2_2
c263.DEAF1/2 V_DEAF1_02
c135.KLF/Sp4 Sp4_secondary
c167.PAX4/3 V_PAX4_01
c183.ARNT::AHR V_AHR_Q5
c128.ARID3A Gata6_secondary
c258.IRF4 Irf6_secondary
c16.CTCF_Core V_CTCF_01
c151.ZNF V_SP1SP3_Q4
c238.HIC1 V_HIC1_02
c34.NFYA V_NFY_01
c208.PAX5/2 MA0014.1–Pax5
c194.GMEB1 Gmeb1_secondary
c197.HSFY2 HSFY2_HSF_2
c116.GLI ZBTB7A_C2H2_1
c192.MYF6 Myf6_secondary
c175.CENPB CENPB_CENPB_1
c58.GMEB2 GMEB2_SAND_2
c219.ZFX V_ZFX_01
c221.GCM1/3 Plagl1_primary
c177.Zic Zic1_primary
c138.CTCF_Upstream CTCF_Upstream
c182.WHN V_WHN_B
c37.MYB MA0100.1–Myb
c12.CTCF_Upstream CTCF_UpstreamP1
Placental
Lymphoid
Myeloid / erythroid
Cardiac
Musculoskeletal
Vascular / endothelial
Embryonic / primitive
Neural
Digestive
Stromal A
Stromal B
Renal
Cancer/ epithelial
Pulmonary devel.
Renal / organ devel.
Tissue invariant
b
GCM1
IRF
ETS/SPI1
MEF2
E-box
ETS/ERG
Oct-4
NeuroD/Olig
HNF4A
AP-1
HNF1
TP63
FOX
PAX2
NRF1
CTCF
CNOT3

### ED Figure 5

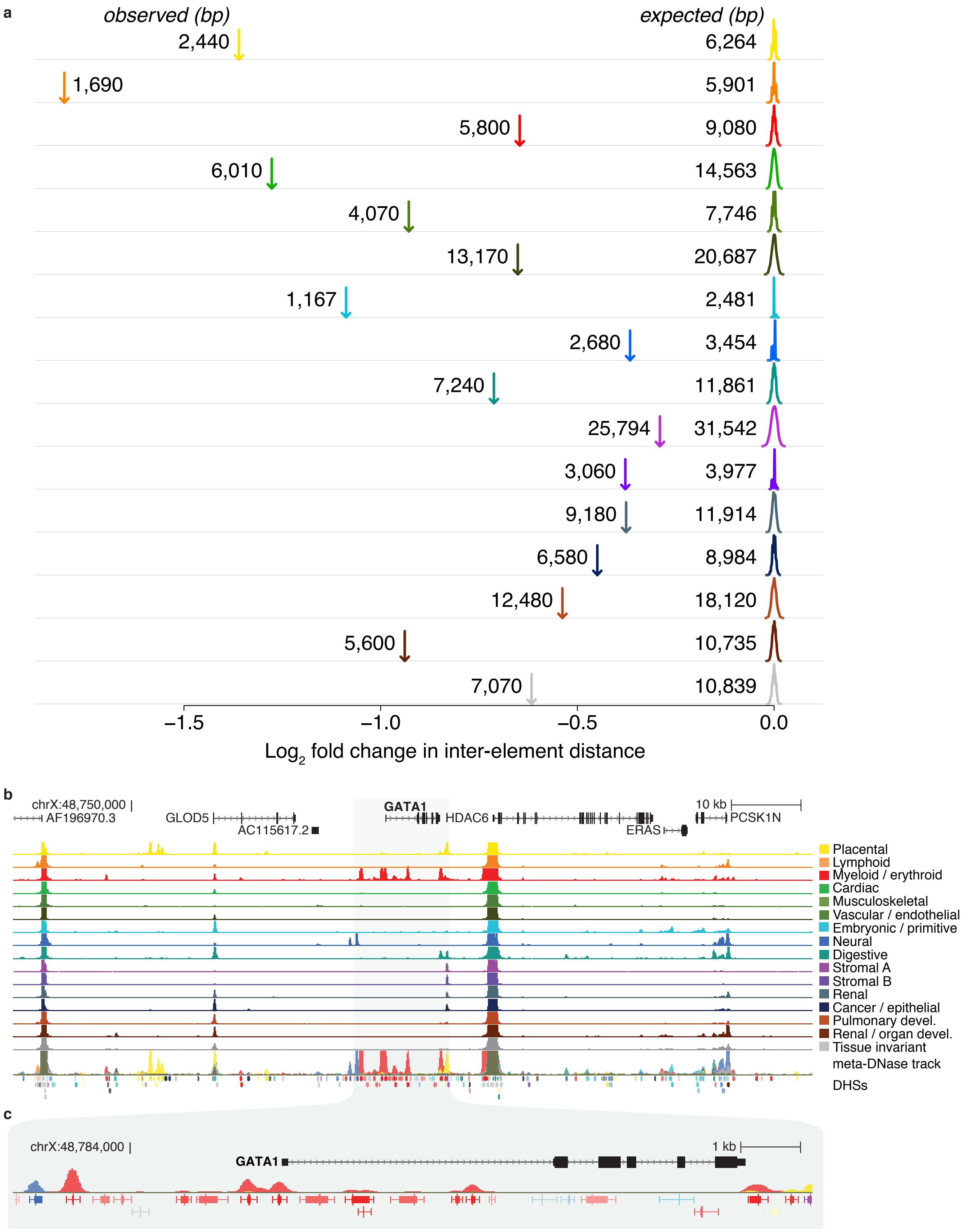
a
observed (bp)
expected (bp)
2,440
6,264
1,690
5,901
5,800
9,080
6,010
14,563
4,070
7,746
13,170
20,687
1,167
2,481
2,680
3,454
7,240
11,861
25,794
31,542
3,060
3,977
9,180
11,914
6,580
8,984
12,480
18,120
5,600
10,735
7,070
10,839
−1.5
−1.0
−0.5
0.0
Log2 fold change in inter-element distance
b
chrX:48,750,000
AF196970.3
GLOD5
AC115617.2
GATA1
HDAC6
ERAS
10 kb
PCSK1N
Placental
Lymphoid
Myeloid / erythroid
Cardiac
Musculoskeletal
Vascular / endothelial
Embryonic / primitive
Neural
Digestive
Stromal A
Stromal B
Renal
Cancer / epithelial
Pulmonary devel.
Renal / organ devel.
Tissue invariant
meta-DNase track
DHSs
c
chrX:48,784,000
1 kb
GATA1

ED Figure 6

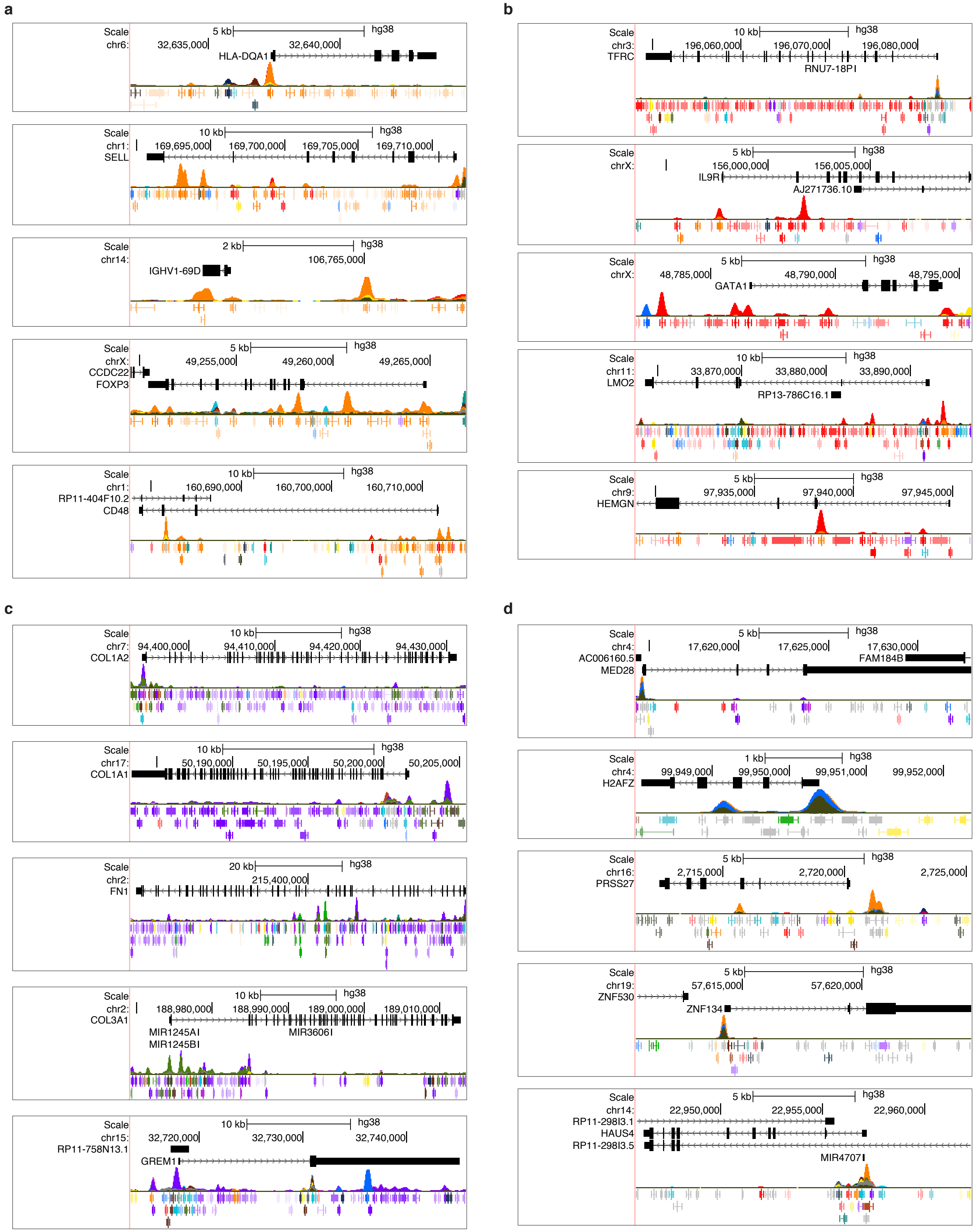

a
Scale
chr6:
5 kb
hg38
32,635,000
32,640,000
HLA-DQA1
Scale
chr1:
SELL
10 kb
hg38
169,695,000
169,700,000
169,705,000
169,710,000
Scale
chr14:
2 kb
hg38
106,765,000
IGHV1-69D
Scale
chrX:
CCDC22
FOXP3
5 kb
hg38
49,255,000
49,260,000
49,265,000
Scale
chr1:
RP11-404F10.2
CD48
10 kb
hg38
160,690,000
160,700,000
160,710,000
b
Scale
chr3:
TFRC
10 kb
hg38
196,060,000
196,070,000
196,080,000
RNU7-18P
Scale
chrX:
5 kb
hg38
156,000,000
156,005,000
IL9R
AJ271736.10
Scale
chrX:
5 kb
hg38
48,785,000
48,790,000
48,795,000
GATA1
Scale
chr11:
LMO2
10 kb
hg38
33,870,000
33,880,000
33,890,000
RP13-786C16.1
Scale
chr9:
HEMGN
5 kb
hg38
97,935,000
97,940,000
97,945,000
c
Scale
chr7:
COL1A2
10 kb
hg38
94,400,000
94,410,000
94,420,000
94,430,000
Scale
chr17:
COL1A1
10 kb
hg38
50,190,000
50,195,000
50,200,000
50,205,000
Scale
chr2:
FN1
20 kb
hg38
215,400,000
Scale
chr2:
COL3A1
10 kb
hg38
188,980,000
188,990,000
189,000,000
189,010,000
MIR1245A
MIR1245B
MIR3606
Scale
chr15:
RP11-758N13.1
GREM1
10 kb
hg38
32,720,000
32,730,000
32,740,000
d
Scale
chr4:
AC006160.5
MED28
5 kb
hg38
17,620,000
17,625,000
17,630,000
FAM184B
Scale
chr4:
H2AFZ
1 kb
hg38
99,949,000
99,950,000
99,951,000
99,952,000
Scale
chr16:
PRSS27
5 kb
hg38
2,715,000
2,720,000
2,725,000
Scale
chr19:
ZNF530
5 kb
hg38
57,615,000
57,620,000
ZNF134
Scale
chr14:
RP11-298I3.1
HAUS4
RP11-298I3.5
5 kb
hg38
22,950,000
22,955,000
22,960,000
MIR4707


ED Figure 7

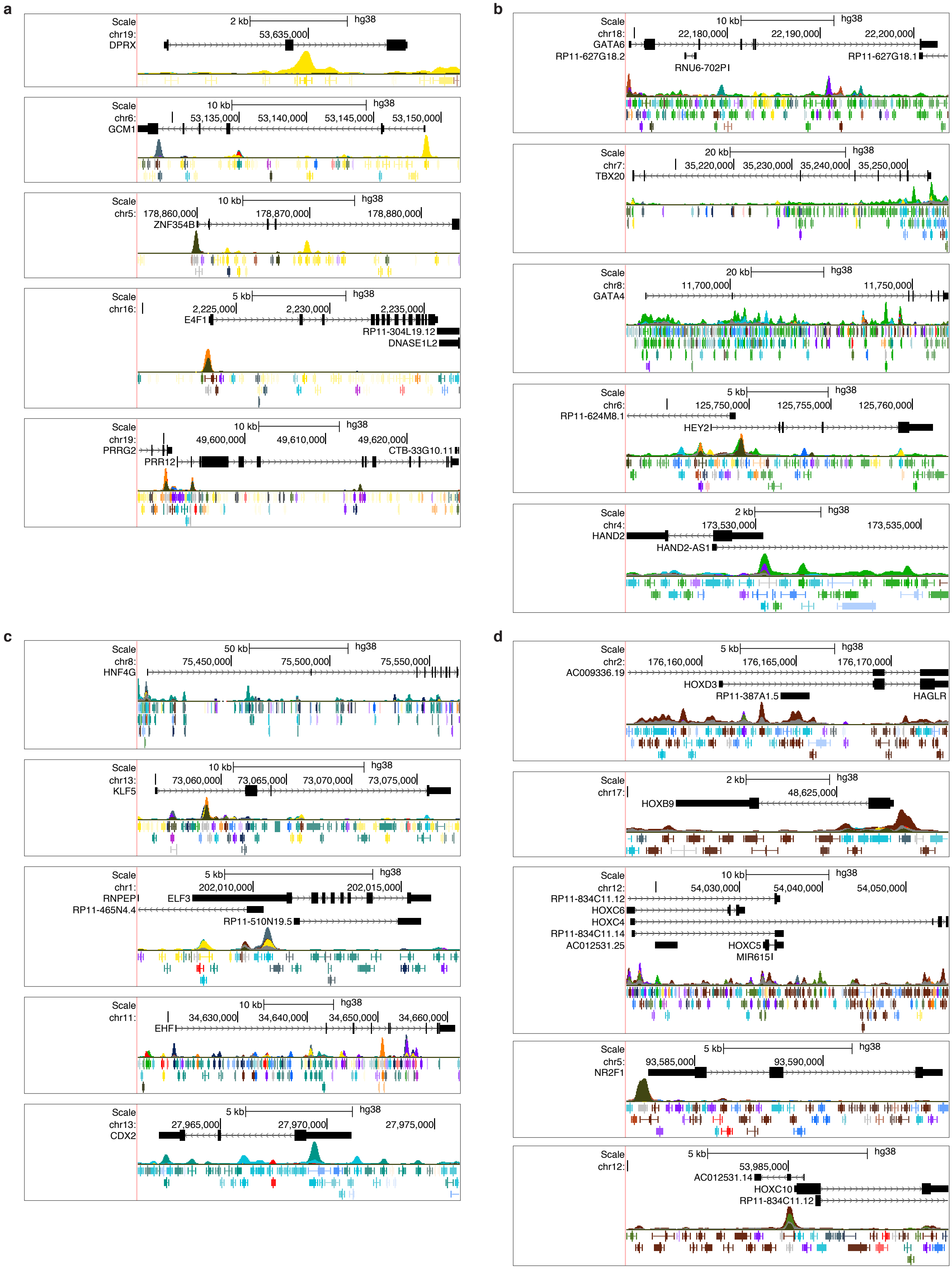
a
Scale
chr19:
DPRX
2 kb
hg38
53,635,000
chr6:
GCM1
10 kb
53,135,000
53,140,000
53,145,000
53,150,000
chr5:
ZNF354B
178,860,000
178,870,000
178,880,000
chr16:
E4F1
RP11-304L19.12
DNASE1L2
5 kb
2,225,000
2,230,000
2,235,000
chr19:
PRRG2
PRR12
CTB-33G10.11
49,600,000
49,610,000
49,620,000
b
chr18:
GATA6
RP11-627G18.2
RNU6-702P
RP11-627G18.1
22,180,000
22,190,000
22,200,000
chr7:
TBX20
20 kb
35,220,000
35,230,000
35,240,000
35,250,000
chr8:
GATA4
11,700,000
11,750,000
chr6:
RP11-624M8.1
HEY2
125,750,000
125,755,000
125,760,000
chr4:
HAND2
HAND2-AS1
173,530,000
173,535,000
c
chr8:
HNF4G
50 kb
75,450,000
75,500,000
75,550,000
chr13:
KLF5
73,060,000
73,065,000
73,070,000
73,075,000
chr1:
RNPEP
ELF3
RP11-465N4.4
RP11-510N19.5
202,010,000
202,015,000
chr11:
EHF
34,630,000
34,640,000
34,650,000
34,660,000
chr13:
CDX2
27,965,000
27,970,000
27,975,000
d
chr2:
AC009336.19
HOXD3
RP11-387A1.5
HAGLR
176,160,000
176,165,000
176,170,000
chr17:
HOXB9
48,625,000
chr12:
RP11-834C11.12
HOXC6
HOXC4
RP11-834C11.14
AC012531.25
HOXC5
MIR615
54,030,000
54,040,000
54,050,000
chr5:
NR2F1
93,585,000
93,590,000
chr12:
AC012531.14
HOXC10
RP11-834C11.12
53,985,000

ED Figure 8

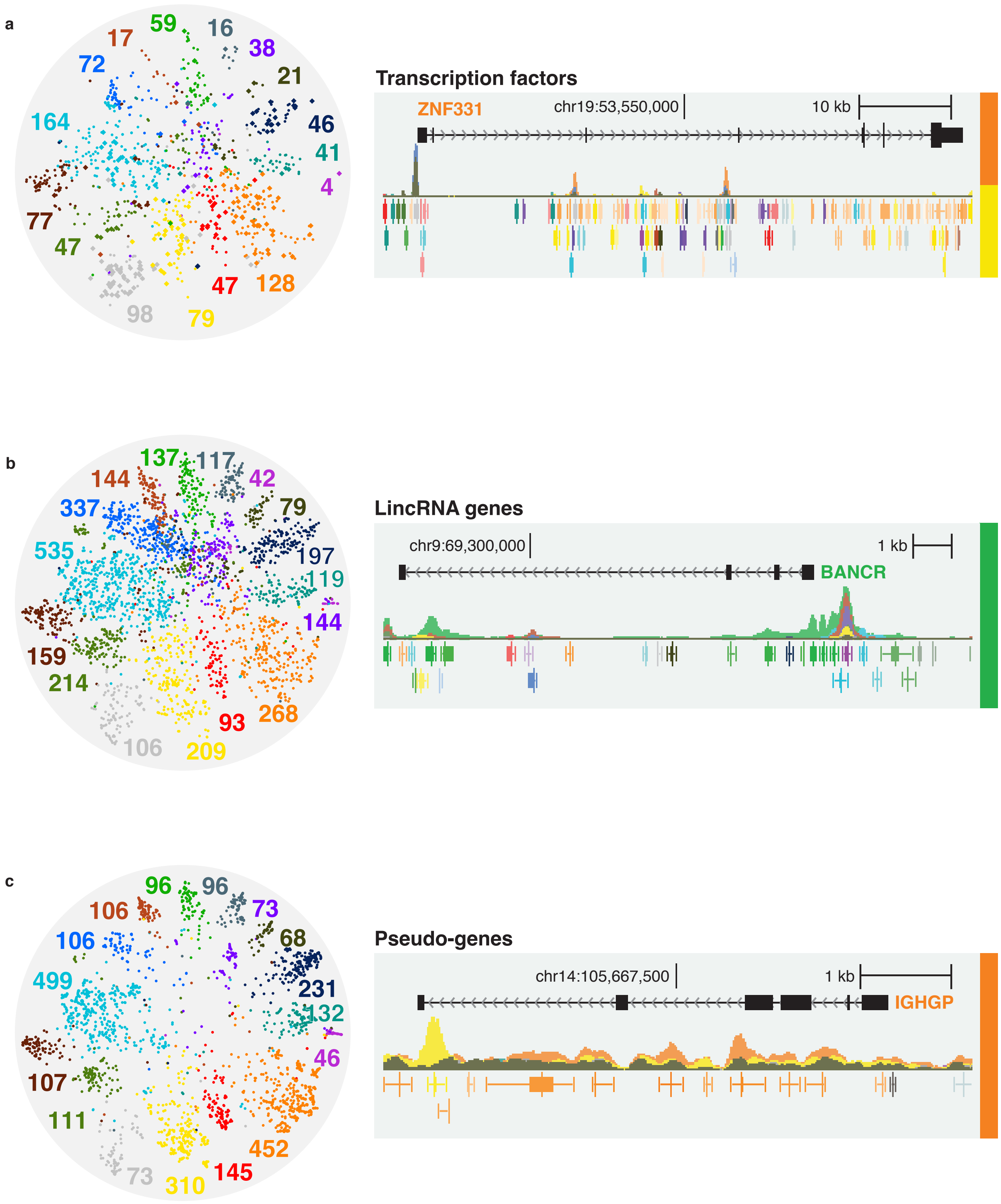
a
17
59
16
38
72
21
164
46
41
4
77
47
47
128
98
79
Transcription factors
ZNF331
chr19:53,550,000
10 kb
b
137
117
144
42
337
79
535
197
119
144
159
214
106
209
93
268
LincRNA genes
chr9:69,300,000
1 kb
BANCR
c
96
96
106
73
106
68
499
231
132
46
107
111
73
310
145
452
Pseudo-genes
chr14:105,667,500
1 kb
IGHGP

ED Figure 9

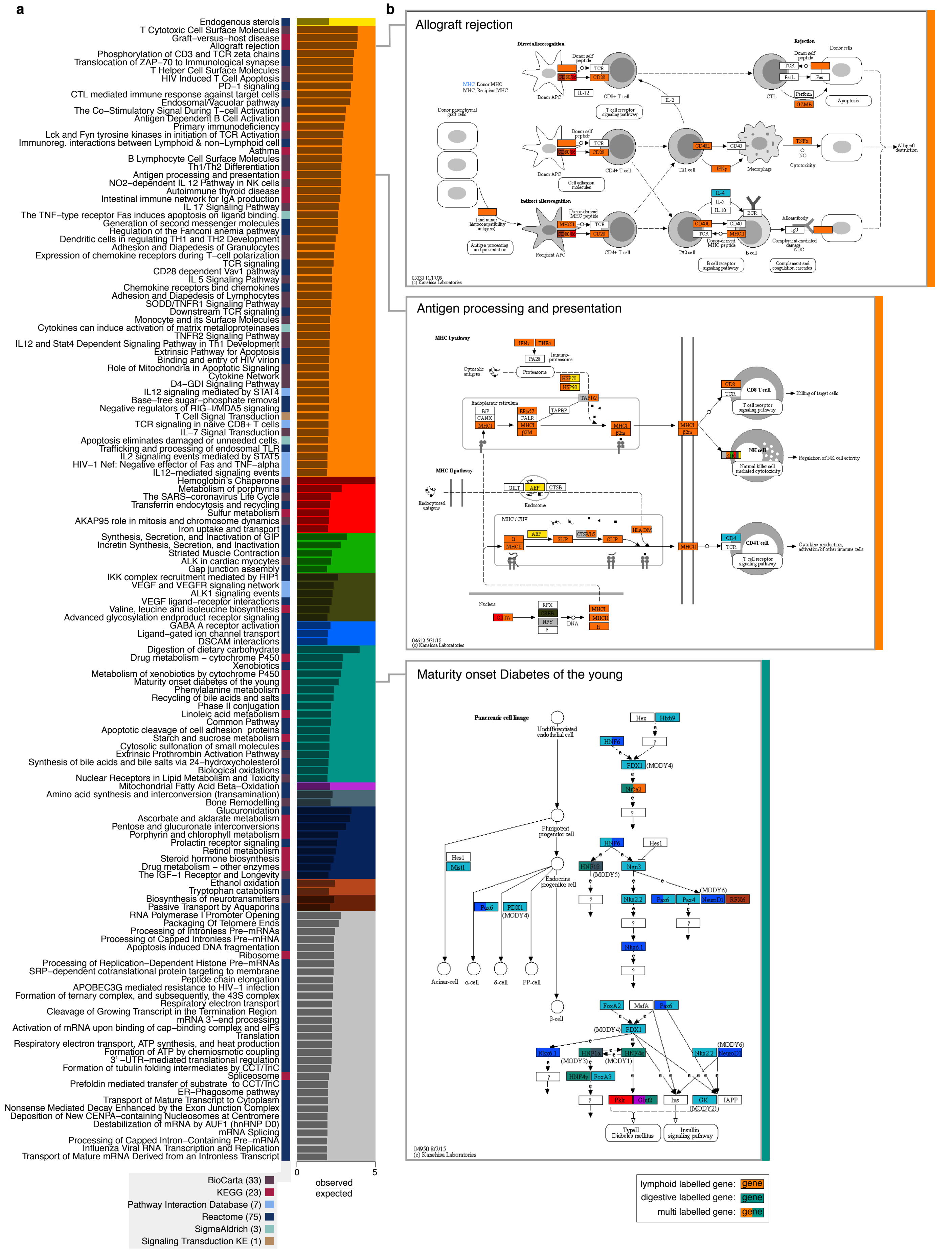
a
Endogenous sterols
T Cytotoxic Cell Surface Molecules
Graft−versus−host disease
Allograft rejection
Phosphorylation of CD3 and TCR zeta chains
Translocation of ZAP−70 to Immunological synapse
T Helper Cell Surface Molecules
HIV Induced T Cell Apoptosis
PD−1 signaling
CTL mediated immune response against target cells
Endosomal/Vacuolar pathway
The Co−Stimulatory Signal During T−cell Activation
Antigen Dependent B Cell Activation
Primary immunodeficiency
Lck and Fyn tyrosine kinases in initiation of TCR Activation
Immunoreg. interactions between Lymphoid & non−Lymphoid cell
Asthma
B Lymphocyte Cell Surface Molecules
Th1/Th2 Differentiation
Antigen processing and presentation
NO2−dependent IL 12 Pathway in NK cells
Autoimmune thyroid disease
Intestinal immune network for IgA production
IL 17 Signaling Pathway
The TNF−type receptor Fas induces apoptosis on ligand binding.
Generation of second messenger molecules
Regulation of the Fanconi anemia pathway
Dendritic cells in regulating TH1 and TH2 Development
Adhesion and Diapedesis of Granulocytes
Expression of chemokine receptors during T−cell polarization
TCR signaling
CD28 dependent Vav1 pathway
IL 5 Signaling Pathway
Chemokine receptors bind chemokines
Adhesion and Diapedesis of Lymphocytes
SODD/TNFR1 Signaling Pathway
Downstream TCR signaling
Monocyte and its Surface Molecules
Cytokines can induce activation of matrix metalloproteinases
TNFR2 Signaling Pathway
IL12 and Stat4 Dependent Signaling Pathway in Th1 Development
Extrinsic Pathway for Apoptosis
Binding and entry of HIV virion
Role of Mitochondria in Apoptotic Signaling
Cytokine Network
D4−GDI Signaling Pathway
IL12 signaling mediated by STAT4
Base−free sugar−phosphate removal
Negative regulators of RIG−I/MDA5 signaling
T Cell Signal Transduction
TCR signaling in naive CD8+ T cells
IL−7 Signal Transduction
Apoptosis eliminates damaged or unneeded cells.
Trafficking and processing of endosomal TLR
IL2 signaling events mediated by STAT5
HIV−1 Nef: Negative effector of Fas and TNF−alpha
IL12−mediated signaling events
Hemoglobin's Chaperone
Metabolism of porphyrins
The SARS−coronavirus Life Cycle
Transferrin endocytosis and recycling
Sulfur metabolism
AKAP95 role in mitosis and chromosome dynamics
Iron uptake and transport
Synthesis, Secretion, and Inactivation of GIP
Incretin Synthesis, Secretion, and Inactivation
Striated Muscle Contraction
ALK in cardiac myocytes
Gap junction assembly
IKK complex recruitment mediated by RIP1
VEGF and VEGFR signaling network
ALK1 signaling events
VEGF ligand−receptor interactions
Valine, leucine and isoleucine biosynthesis
Advanced glycosylation endproduct receptor signaling
GABA A receptor activation
Ligand−gated ion channel transport
DSCAM interactions
Digestion of dietary carbohydrate
Drug metabolism − cytochrome P450
Xenobiotics
Metabolism of xenobiotics by cytochrome P450
Maturity onset diabetes of the young
Phenylalanine metabolism
Recycling of bile acids and salts
Phase II conjugation
Linoleic acid metabolism
Common Pathway
Apoptotic cleavage of cell adhesion proteins
Starch and sucrose metabolism
Cytosolic sulfonation of small molecules
Extrinsic Prothrombin Activation Pathway
Synthesis of bile acids and bile salts via 24−hydroxycholesterol
Biological oxidations
Nuclear Receptors in Lipid Metabolism and Toxicity
Mitochondrial Fatty Acid Beta−Oxidation
Amino acid synthesis and interconversion (transamination)
Bone Remodelling
Glucuronidation
Ascorbate and aldarate metabolism
Pentose and glucuronate interconversions
Porphyrin and chlorophyll metabolism
Prolactin receptor signaling
Retinol metabolism
Steroid hormone biosynthesis
Drug metabolism − other enzymes
The IGF−1 Receptor and Longevity
Ethanol oxidation
Tryptophan catabolism
Biosynthesis of neurotransmitters
Passive Transport by Aquaporins
RNA Polymerase I Promoter Opening
Packaging Of Telomere Ends
Processing of Intronless Pre−mRNAs
Processing of Capped Intronless Pre−mRNA
Apoptosis induced DNA fragmentation
Ribosome
Processing of Replication−Dependent Histone Pre−mRNAs
SRP−dependent cotranslational protein targeting to membrane
Peptide chain elongation
APOBEC3G mediated resistance to HIV−1 infection
Formation of ternary complex, and subsequently, the 43S complex
Respiratory electron transport
Cleavage of Growing Transcript in the Termination Region
mRNA 3'−end processing
Activation of mRNA upon binding of cap−binding complex and eIFs
Translation
Respiratory electron transport, ATP synthesis, and heat production
Formation of ATP by chemiosmotic coupling
3' −UTR−mediated translational regulation
Formation of tubulin folding intermediates by CCT/TriC
Spliceosome
Prefoldin mediated transfer of substrate to CCT/TriC
ER−Phagosome pathway
Transport of Mature Transcript to Cytoplasm
Nonsense Mediated Decay Enhanced by the Exon Junction Complex
Deposition of New CENPA−containing Nucleosomes at Centromere
Destabilization of mRNA by AUF1 (hnRNP D0)
mRNA Splicing
Processing of Capped Intron−Containing Pre−mRNA
Influenza Viral RNA Transcription and Replication
Transport of Mature mRNA Derived from an Intronless Transcript
0
5
observed
expected
BioCarta (33)
KEGG (23)
Pathway Interaction Database (7)
Reactome (75)
SigmaAldrich (3)
Signaling Transduction KE (1)
b
Allograft rejection
05330 11/17/09
(c) Kanehisa Laboratories
Antigen processing and presentation
04612 5/31/18
(c) Kanehisa Laboratories
Maturity onset Diabetes of the young
04950 8/7/15
(c) Kanehisa Laboratories
lymphoid labelled gene: gene
digestive labelled gene: gene
multi labelled gene: gene

ED Figure 10

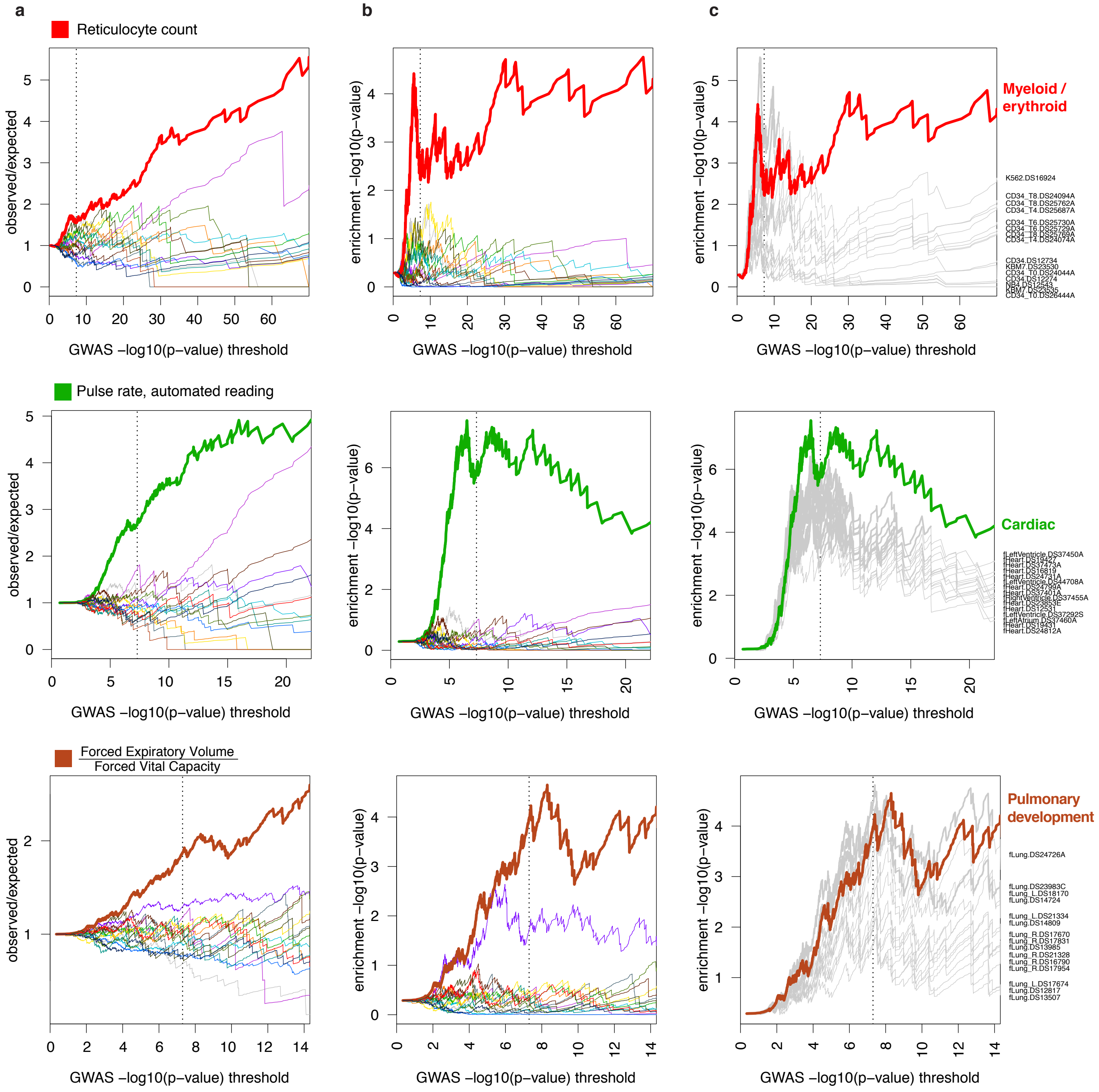
a
b
c
Reticulocyte count
Pulse rate, automated reading
Forced Expiratory Volume
Forced Vital Capacity
observed/expected
enrichment −log10(p−value)
GWAS −log10(p−value) threshold
Myeloid / erythroid
Cardiac
Pulmonary development

ED Figure 11

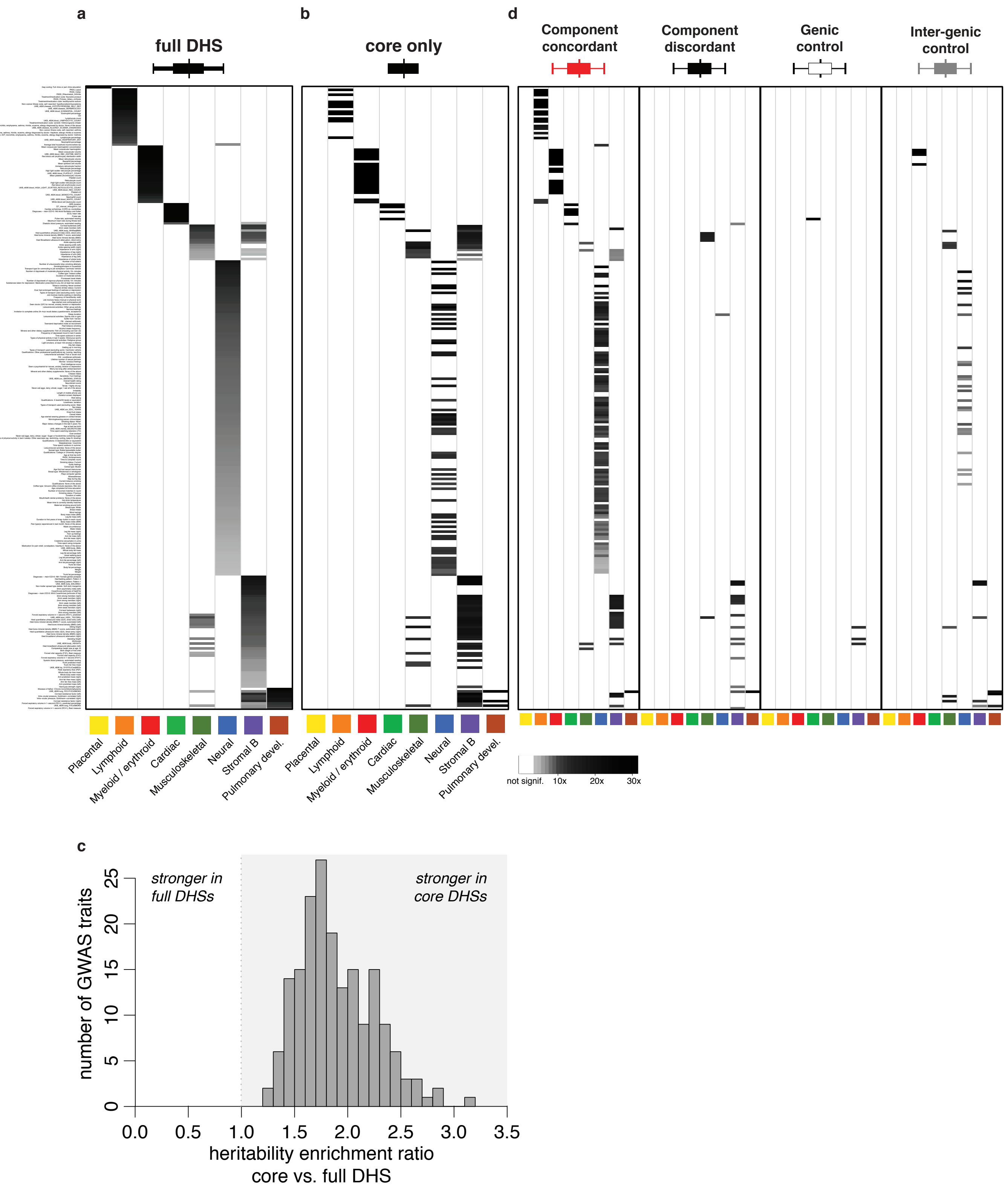
a
full DHS
b
core only
d
Component concordant
Component discordant
Genic control
Inter-genic control
Placental
Lymphoid
Myeloid / erythroid
Cardiac
Musculoskeletal
Neural
Stromal B
Pulmonary devel.
not signif.
10x
20x
30x
c
stronger in full DHSs
stronger in core DHSs
number of GWAS traits
0
5
10
15
20
25
0.0
0.5
1.0
1.5
2.0
2.5
3.0
3.5
heritability enrichment ratio core vs. full DHS

ED Figure 12
