## Supplemental Legends for "Index and biological spectrum of accessible DNA elements in the human genome"

### Extended Data Figure Legends

**Extended Data Figure 1 — Construction of a DHS index**

**a-b.** Delineation of index DNase I Hypersensitive Sites (DHSs) from raw DNase-seq signal tracks, shown for simplified data (**a**) and actual data (**b**). Starting from individual DNase-seq datasets (step 1), we call peaks in each dataset (step 2), aggregate peak summits into clusters, indicating isolated accessibility events (step 3), group full peak coordinates according to these clusters (step 4), and delineate DHSs using Full-Width at Half Maximum (FWHM) (step 5). **c.** Detailed view of FWHM delineation. **d-e.** Confidence scores based on DNase I signal strengths assigned to each DHS, allowing for pragmatic filtering using either summed (**d**) or mean (**e**) signal strength — the former assigning high confidence scores to DHSs with overall high signal levels across datasets, the latter providing a score normalized by the number of datasets in which a DHS was observed.

**Extended Data Figure 2 — Genomic context of DHS index elements**

**a.** Overall coverage of 3.5M+ DHSs across genes and repetitive elements. **b.** Coverage of classes and families of repetitive elements. **c.** Coverage of annotated genic regions. **d.** Density plot of DHS distance to their closest transcription start site (TSS), showing that the vast majority of DHSs are found distal to annotated promoters. **e.** DHS summits show an increase in sequence conservation (phyloP) and a decrease in within-human sequence variation (TOPMed, $\pi \times 10^4$). **f.** Histogram indicating the variety in cell type selectivity of DHSs, ranging from single cell types to groups of 10s, 100s or even all assayed cellular conditions.

**Extended Data Figure 3 — NMF decomposition of DHS index**

**a**. Schematic depiction of Non-Negative Matrix Factorization (NMF) applied to an *n*-by-*m* matrix resulting in *k* components. The objective is to minimize the difference between the original matrix **V** and the product of **W** and **H**, such that all elements of **W** and **H** are non-negative. **b**. Depiction of NMF applied to our DNase-seq dataset of 733 biosample datasets and 3.5M+ DHSs, using *k* components. **c**. Color-based view from the values shown in **b**. Colors indicate relative loadings of each NMF component, for both biosamples and DHSs. **d.** Choice of NMF decision boundary (0.35) based on maximal F1 score as a function of number of components *k* (4 to 36). **e.** F1 score as a function of the number of components *k*, with the chosen *k*=16 and corresponding F1 score indicated. **f.** Gradient showing reduced gain in F1 score after *k*=16.

**Extended Data Figure 4 — Association of regulatory components with cellular conditions**

**a.** Barplots showing for each NMF component the top 15 DNase-seq datasets in terms of NMF loadings. NMF loading strength (x-axes) and dataset labeling (y-axes) are indicated. **b.** Boxplots showing for each NMF component its loadings across those biosamples for which that component is maximally loaded. **c.** Beyond the top 15 biosamples for each component, general associations of components with annotations regarding human organ systems and cancer. Indicated are Bonferroni corrected p-values, resulting from Mann-Whitney U tests. **d.** Distribution of biosamples across (maximal) NMF components, for the number of components (*k*) ranging from 2 to 16. Labels at the top indicate at which point distinct lineages became represented in corresponding components.

**Extended Data Figure 5 — Association of regulatory components with TF motifs**
**a.** Enrichment of transcription factor (TF) binding motifs in regulatory components. Grayscale values indicate enrichment levels, and only statistically significant results are included. Regulatory components are shown on the x-axis, and TF motifs (clusters) on the y-axis. Shown are a maximum of one motif per motif cluster (see Methods). **b.** Selected top-enriched TF motifs for each regulatory component.

**Extended Data Figure 6 — Regional clustering of same-component DHSs around genes**
**a.** Component-specific genomic clustering of DHSs, as shown by the median distance between same-component DHSs as compared to the median distance after random permutation of DHS-component labels. **b.** Regulatory landscape +/- 50kb around the GATA1 gene, indicating GENCODE v28 gene annotations, meta-DNase tracks for individual regulatory components (see Methods), and a meta-DNase overlay track. **c.** Detailed view, restricted to the GATA1 regulatory landscape, including its delineated and annotated DHSs. Collectively, this landscape shows a statistical over-representation of DHSs associated with the myeloid/erythroid (red) component.

**Extended Data Figure 7 — Top 5 labeled protein-coding genes for selected components**
Top-scoring protein-coding genes per regulatory component reflect their functional roles, as shown for lymphoid (**a**), myeloid / erythroid (**b**), stromal (**c**) and tissue-invariant (**d**) components. For each gene, the full extent of its regulatory landscape used for labeling is shown, including GENCODE v28 gene annotations, meta-DNase overlay track, and annotated DHSs.

**Extended Data Figure 8 — Top 5 labeled TF genes for selected components**
Top-scoring transcription factor (TF) genes per regulatory component reflect their functional roles, as shown for placental (**a**), cardiac (**b**), digestive (**c**) and organ developmental / renal (**d**) components. For each gene, the full extent of its regulatory landscape used for labeling is shown, including GENCODE v28 gene annotations, meta-DNase overlay track, and annotated DHSs.

**Extended Data Figure 9 — 2D projections of enrichment patterns for gene categories**
2D projection coordinates generated using t-SNE on all genes significantly associated with a regulatory component and shown selectively for subsets of gene categories, namely transcription factors (**a**), lincRNA genes (**b**) and pseudo-genes (**c**). Indicated are the number of labelled genes in each combination of gene category and regulatory component. Examples of labeled genes are shown as follows. **a.** Regulatory landscape of ZNF331; a poorly annotated zinc-finger (ZNF) TF, annotated with the lymphoid and placental components. **b.** Regulatory landscape of BANCR; a long intergenic non-coding RNA (lincRNA) gene, recently associated with cardiomyocyte migration. **c.** Regulatory landscape of IGHGP; a pseudo-gene associated with the lymphoid component.

**Extended Data Figure 10 — Component-wise labeling of pathways**
Labeling of MSigDB canonical pathways using regulatory components, through the regulatory landscapes of constituent genes. **a.** Pathways with a significant association (FDR < 5%) and an observed/expected ratio of at least 2. The most strongly associated components and their observed/expected ratios are indicated for each pathway, as well as their source databases. **b.** Examples of three component-associated pathways from the KEGG database, with genes colored according to their (significant) majority component(s).

**Extended Data Figure 11 — Quantitative association of components with GWAS traits**
**a-c.** Quantitative association of component-DHSs with GWAS traits reticulocyte count, pulse rate, and FEV1/FVC ratio. The canonical genome-wide significance threshold ($5 \times 10^{-8}$) is indicated. **a.** Enrichment ratio for each regulatory component as a function of the set of variants with a minimal indicated significance stringency threshold. **b.** Enrichment $-\log_{10}$(p-value) for each regulatory component. **c.** Exclusively showing enrichment $-\log_{10}$(p-value) for the single strongest regulatory component, along with its most strongly associated biosamples.

**Extended Data Figure 12 — GWAS trait associations of regulatory components**
**a-b**. GWAS trait associations with index DHSs for regulatory components. Grayscale values indicate heritability enrichment levels for statistically significant traits. Shown are heritability enrichment results based on the full delineated width of index DHSs (**a**) and restricted to index DHS 'core' regions (**b**). **c.** Ratio of heritability enrichment values for 'core' versus 'full size' DHSs. **d.** Heritability enrichments stratified according to gene landscape DHS types.

### Supplementary Table Legends

**Supplementary Table 1 — Biosample metadata spreadsheet**

Spreadsheet with per-biosample metadata, including biosample taxonomy and identifiers, experimental and computational quality measures of resulting datasets, and hyperlinks to experimental protocols.

**Supplementary Table 2 — Gene labeling spreadsheet**

Spreadsheet with per-component gene labelings.
