## Supplemental Methods for "Index and biological spectrum of accessible DNA elements in the human genome"

**Methods** **2**
Primary data generation and processing 2
Generation of DNase I hypersensitivity maps 2
DNase-seq sample selection and data processing 2
**Construction of an Index of DNA accessibility** **3**
Detection of strong peaks in individual datasets 3
Definition of isolated accessibility events 3
Delineation of consensus DHSs 3
Annotation of individual DHSs 4
Overlap of the DHS Index with genomic annotations 5
**Construction of a DHS Vocabulary** **5**
Rationale for component-wise description of DHSs 5
Decomposition using Non-negative Matrix Factorization (NMF) 6
NMF implementation and choice of number of components 6
Labeling of NMF components and DHSs 7
**Regulatory annotation of human genes** **8**
Per-component genomic distribution of DHSs 8
Per-component meta-DNase tracks 8
Definition of regulatory landscapes and component associations 9
Annotations for GENCODE genes and pseudo-gene types 9
Visualization of gene regulatory annotations 9
Construction and use of gene expression compendium dataset 9
Annotation and visualization of pathway labellings 10
**Genetic variation analyses** **10**
GWAS traits and summary statistics 10
Estimates of SNP-based heritability 10
Quantitative trait associations 11
Stratified LD-score regression 11
Heritability enrichments for DHS Vocabulary components 11
Unique per-annotation contributions to SNP-based heritability 12
**Data and Code Availability** **12**
Biosample metadata 12
Index of consensus DHSs 12
Construction of DHS Vocabulary 12
**Supplemental References** **13**

### Methods

#### Primary data generation and processing

##### Generation of DNase I hypersensitivity maps

DNase I assays were generally performed according to a protocol detailed previously[1]. This protocol involves treatment of intact nuclei with the small enzyme DNase I which is able to penetrate the nuclear pore and cleave exposed DNA. Small (< 1kb) fragments are isolated from lysed nuclei following DNase I treatment, linkers are added, and the resulting library is sequenced. Because tissue and cell culture and isolation and handling protocols differ for different biosamples, these are all available online and indexed with URLs in **Supplementary Table 1** (Google Spreadsheet).

##### DNase-seq sample selection and data processing

Because regulatory DNA accessibility exhibits a wide dynamic range (hundreds-fold) across elements, the full extent of the accessible DNA landscape of a given cell type only becomes evident with high quality, high signal-to-noise data. At the high end of the spectrum are promoter elements, which exhibit high accessibility but little cell type-selectivity. By contrast, cell selectivity is concentrated in moderate-to-lower accessibility distal elements, which are under-sampled in lower quality data.

For this reason, we considered exclusively high quality data with a SPOT score of at least 0.3 for our analyses, yielding a total of 733 DNase-seq datasets. Collectively, these data assay a total of 439 cell and tissue states spanning all human organ systems. We defined the term "cell and tissue state" to encompass distinct cell types; distinct developmental time-points in case of fetal tissue data; different time-points in case of differentiating cultured cells; and (for a small number of datasets) different treatment conditions (**Supplementary Table 1;** Google Spreadsheet).

Data processing of DNase-seq FASTQ files was done using the ENCODE DCC DNase-HS pipeline, version 2, paired-end (ENCPL202DNS). Briefly, reads were trimmed of adapter match sequence and subsequently aligned to the human genome reference GRCh38/hg38 (ENCSR425FOI) using BWA (version 0.7.12). Filtered, quality mapped, reads from all sequencing runs of a given library were merged. Collectively across datasets, we obtained 94,091,469,219 (94 billion) usable mapped DNase I cleavage fragments.

Further processing and QC evaluation was performed in accordance with the ENCODE DCC pipeline specification (https://www.encodeproject.org/data-standards/dnase-seq/). Standardized output from the DCC pipeline includes normalized read densities, which are a sum of 5' read end counts in a 150bp sliding window across the genome. These values are reported at 20bp steps across the genome. In order to facilitate integrative analyses across datasets, these values are normalized by dividing each value by the total uniquely mapping read count and scaling by 1 million. Non-normalized and non-windowed versions of these read densities are also output by the DCC pipeline. These processed DNase-seq datasets are available from the ENCODE Consortium Data Portal (http://www.encodeproject.org).

#### Construction of an Index of DNA accessibility

For any given DNase I Hypersensitive Site (DHS) shared across a number of datasets, the precise observed location and width of said hypersensitive site may vary slightly between datasets. For this reason, an integrative analysis across multiple datasets requires a form of alignment of sites. In essence, each DNase-seq peak call represents an imperfect guess of where the actual hypersensitive site is. We exploited this behavior to obtain more high-resolution and precisely delineated lists of regulatory regions by combining evidence across datasets to arrive at a consensus delineation. The code for this procedure is available (https://github.com/Altius/Index) and a brief description follows.

##### Detection of strong peaks in individual datasets

Starting from per-dataset data tracks (**Extended Data Fig. 1a-b, step 1**), we detected statistically strong (FDR < 0.1%) variable-width peaks across individual datasets using the Hotspot2 software (https://github.com/Altius/hotspot2). Hotspot2 leverages an enzymatic DNA cleavage model to measure enrichment over background and identify regions of cleavage enrichment, along with accompanying confidence values. Alignment files were input into the pipeline and processed over uniquely mappable regions of the genome. On average, each dataset was called with 104,433 variable-width peaks (FDR < 0.1%), for a total of 76,549,656 (76.5M) peaks across all 733 DNase-seq datasets (**Extended Data Fig. 1a-b, step 2**). For each peak, we registered their start, end and estimated summit positions, as well as their signal intensities.

##### Definition of isolated accessibility events

We used the local clustering of peak summit positions across datasets to define isolated events of chromatin accessibility. More specifically, we aggregated the 1bp summit locations of peaks across all datasets, as described next (**Extended Data Fig. 1a-b, step 3**). For computational efficiency, we parallelized our procedure by dividing up the genome into roughly 5000 inter-independent regions, separated by large regions containing zero peak summits (see code for details). We further subdivided regions of subsequent peak summits if separated more than 20bp, the resolution of our base signal, or if the number of peak summits dropped below the median number of peak summits in that region (with a minimum of 1 per base-pair), allowing for the recovery of substructures within tightly clustered DNase I hypersensitive sites. This resulted in a tentative set of isolated accessibility events, anchored on sets of peak summits (**Extended Data Fig. 1a-b, step 4**).

##### Delineation of consensus DHSs

We assigned genomic coordinates to each isolated event of chromatin accessibility by accumulating contributing peak positions (full start-to-end coordinates) and delineating the Full-Width at Half Maximum (FWHM) on the resulting signal, representing a consensus across datasets (**Extended Data Fig. 1a-b, step 5**). Starting from local maxima of summit positions, we subsequently included neighboring base-pairs in each direction, up until half of the height at the local maxima. This FWHM approach for delineating areas of consistent enrichment results in base pair-level delineations supported by at least half of all datasets in which a peak was called at that location.

Each delineation was assigned a consensus summit and score (see below). Two delineations were combined in case the summit of one delineation overlaps the coordinates of another delineation with a higher score. We combine delineations by merging the two (peak) summit clusters and re-estimating a single FWHM delineation, to retain high resolution consensus delineations. In a few cases, this step was skipped because it would result in a single delineation containing a relatively large number of datasets contributing more than one peak. More specifically, if the result of combining two delineations would be that if the majority of involved datasets contribute more than one peak, the two delineations did not get merged (see code for implementation details).

The above resulted in an Index of 3,591,898 DNase I Hypersensitive Sites (DHSs), each occurring across any number of 733 input DNase-seq datasets. We next annotated each individual DHS.

##### Annotation of individual DHSs

Through the above procedure we obtained perfect provenance of which DNase-seq peaks contribute to each DHS delineation. We used this to estimate not only the coordinates of DHSs (i.e. through FWHM), but also the most likely center-of-mass, or centroid. Specifically, we estimated this centroid position by the median summit position of all contributing peaks. We further defined a 'core' region for each DHS, consisting of the coordinate range containing 95% of all contributing peak summits, i.e. the 2.5% trimmed range. This is conceptually similar to the interquartile range (IQR), except with a breakdown point of 2.5% instead of 25%. It provides us with a measure of dispersion, or positional stability, for each DHS (**Fig. 1d, Extended Data Fig. 1c**). We used this adapted version of the IQR over the standard IQR or confidence interval as the latter two often result in intervals too narrow to meaningfully visualize.

The normalized DNase-seq signal values across datasets provide the basis for per-DHS confidence scores, either summed across all datasets (**Extended Data Fig. 1d**), or by way of the mean signal across summit-contributing datasets (**Extended Data Fig. 1e**). The former quantifies confidence by way of overall signal level across datasets but may penalize dataset-specific or -restricted DHSs. The latter allows more readily for high-scoring dataset-specific DHSs, but does not directly consider the level of replication across datasets. For most practical purposes we recommend the use of confidence scores based on the mean signal across datasets, along with the number of datasets in which a DHS is found.

Lastly, to facilitate an unambiguous identification of Index DHSs, and accommodate posterity of analyses and results, we propose a flexible naming scheme for individual DHSs. Each DHS was assigned a unique identifier, consisting of the chromosome on which it is located, separated by a period with a number reflecting the approximate chromosomal location percentile (**Fig. 1d**). As such, these identifiers read like floating point numbers, allowing for trivial addition of future newly discovered DHSs in-between existing ones. Additionally, because of this, the length of the identifier reflects the approximate local density of DHSs. These features make this identification system relatively robust to future genome builds and portable to personal genomes. The very nature of the actual identifiers makes them interpretable, as opposed to other databases such as dbSNP (e.g. rs1045642) or Ensembl (e.g. ENSG00000210049). Moreover, the system allows for a direct integration with DNase I Footprint IDs built on DHS IDs (chr.DHS_ID.FP_ID, e.g., footprint 11.149576.2).

The resulting set of 3.5M+ DHS delineations is available in tab-separated format from the ENCODE DCC portal (https://www.encodeproject.org/annotations/ENCSR857UZV/) and via Zenodo (https://doi.org/10.5281/zenodo.3542126).

##### Overlap of the DHS Index with genomic annotations

To assess the overlap of our DHS Index elements with repetitive elements (**Extended Data Fig. 2b**), we obtained RepeatMasker[3] annotations downloaded from the University of California Santa Cruz (UCSC) Table Browser[4], and considered the various repeat classes and (sub)families as provided. To perform analogous analyses for human gene annotations (**Extended Data Fig. 2c**), we obtained GENCODE[5] v28 Basic annotations. We defined exons as specified in the GENCODE annotation, promoters as the transcription start site (TSS) of genes +/- 1kb, and introns as the rest of the gene body. Intergenic regions were defined as those not covered by gene bodies or defined promoters. We assigned Index DHSs to these annotations based on maximal overlap.

TOPMed within-human sequence variation data were obtained from the Bravo website (https://bravo.sph.umich.edu/freeze5/hg38/download, Freeze 5, hg38, VCF format). Briefly, we processed the 463M+ variants based on the PASS flag, resulting in a total of 430,235,017 variants used for subsequent analysis. Single-base substitutions were converted to nucleotide diversity scores (π), with a score of zero implied for every genomic base position with no variants. Per-base, phyloP [6] sequence conservation scores were downloaded as-is (http://hgdownload.cse.ucsc.edu/goldenpath/hg38/phyloP100way/). Within-human sequence variation data ($\pi \times 10^4$) and phyloP conservation scores were aligned relative to DHS summits using 20-bp non-overlapping windows tiled across a 1kb region centered on each DHS summit (**Extended Data Fig. 2e**). For each window offset relative to the DHS summit midpoint, genome-wide per-base scores were subsetted using bedops[7] and averaged with GNU datamash.

#### Construction of a DHS Vocabulary

##### Rationale for component-wise description of DHSs

Cluster analysis methods such as hierarchical or k-means clustering have been widely employed to define patterns of gene or regulatory element activity across a range of cell types and states[8,9]. Indeed, hierarchical and partitional clustering has been used to group DHSs with similar behavior across DNase-seq datasets[10,11]. With this approach, each DHS is assigned to a single cluster only, along with other DHSs that display a similar patterning across all datasets. To capture complex patterning of millions of DHSs across >700 datasets with clustering, thousands of clusters would be required, eclipsing utility and interpretability. This requirement arises because different clusters are in effect capturing incomplete snapshots of the operation of what are in reality a much smaller number of co-occurring biological processes. Moreover, these clusters are unlabeled, each consisting of a complex combination of contributions from different cellular conditions. This makes downstream analyses impractical and does not lead to an improved understanding of the repertoire of regulatory patterns present in the data. As such, clustering is poorly suited to capture the rich spectrum of cross-cell-type DHS activation behavior in a biologically interpretable fashion that is robust and broadly useful for both genome annotation and downstream analytical applications.

Beyond clustering, methods have been proposed to learn chromatin state models from genome-wide data[12,13]. Although primarily designed to model data in a temporal fashion (i.e. along a genome), chromatin state models have been trained jointly across large numbers of cell types in a "stacked" manner in an attempt to model complex inter-cell type patterning[14]. As with clustering, for DNase-seq data across many cell types, this approach typically results in a very large number of chromatin states that cannot be directly interpreted across the context of multiple cell types, requiring post-hoc processing to aid in downstream interpretation.

Here, we decompose the occurrence patterns of DHSs across DNase-seq datasets into a small number of components. By combining components, we can describe the behavior of DHSs across multiple cellular conditions, e.g. a DHS occurring in both neural and immune cell types. This way, a DHS occurring in a single cellular condition may be described by a single component, whereas a DHS occurring in many different cellular conditions may be described by a combination of multiple components. By combining even a modest number of components, with varying relative contributions, we are able to capture a vast combinatorial space.

#### Decomposition using Non-negative Matrix Factorization (NMF)

We employed Non-negative Matrix Factorization (NMF)[15,16] to perform the decomposition of the patterning of DHSs across datasets. Originally introduced in the field of computer vision, in bioinformatics research, the major application has so far been to gene expression datasets[17–21]. NMF is distinguished from techniques such as principal component analysis (PCA)[22] or clustering methods that learn holistic rather than parts-based representations of data. NMF aims to model individual 'parts' or 'components' that in combination describe a larger data pattern or object (e.g., nose, eyes, mouth → face), while holistic methods aim to capture global patterns of variation in data (e.g., shadow patterns over a whole face). An additional advantage of NMF over e.g. PCA, is that due to the non-negativity constraint, the learned components can be combined in an interpretable manner, like ingredients in a recipe.

Briefly, NMF aims to break up an $n$-by-$m$ ($n$ datasets, $m$ DHSs) dimensional matrix $\mathbf{V}$ into two new matrices $\mathbf{W}$ (dimensionality $n$-by-$k$) and $\mathbf{H}$ (dimensionality $k$-by-$m$) which can be multiplied together to approximately reconstruct the original data (**Extended Data Fig. 3a-c**). NMF is an unsupervised algorithm that reduces the dimensionality of data when $k$ is less than both $n$ and $m$. In essence, the matrices $\mathbf{W}$ and $\mathbf{H}$ provide a reformulation of DNase-seq datasets and DHSs in terms of $k$ components, instead of in terms of $m$ DHSs or $n$ datasets, respectively.

The defining characteristic of the method is the non-negativity constraint, which means that all elements in $\mathbf{W}$ and $\mathbf{H}$ are constrained to be either zeros or positive values. The non-negativity constraint guarantees that all components are additive, ensuring that the data is represented in a modular fashion. As such, it is an appropriate architecture for chromatin accessibility across large numbers of DNase-seq datasets, where we observe regions of hypersensitivity across all scenarios ranging from being specific to a single dataset to being shared across all datasets.

#### NMF implementation and choice of number of components

Like other dimensionality reduction methods, NMF does not guarantee a total recapitulation of the original data. Instead of searching for an exact reconstruction, we chose to allow information loss in exchange for a more interpretable result. Therefore, we considered using a much smaller number of *k* components than the lower of the two dimensions of our input matrix (733 DNase-seq datasets). To keep the reconstruction error in check, we employed an objective function that is minimized subject to the Frobenius norm (**Extended Data Fig. 3a**). NMF typically uses a random initialization step, leading to unstable results. To alleviate this, we performed the initialization step using singular value decomposition (SVD)[23,24], leading to consistent results while maintaining a performance that is on par with randomly initialized instances. More implementation details follow.

We applied NMF decomposition to a binary matrix **V** consisting of presence/absence calls of *m* DHSs across *n* DNase-seq datasets. The number of components *k* to use is a parameter of the NMF algorithm and its choice is a trade-off between how much of the Biological complexity should be captured versus how interpretable the model should be. To systematically choose a good value for *k*, we assessed the quality of the decomposition across a wide range of values using appropriate metrics as follows.

With only 4% non-zero elements, matrix **V** is quite sparse, and thus we employed metrics suitable for highly class-imbalanced and sparse datasets. We calculated `precision` (TP/(TP+FP)) and `recall` (TP/(TP+FN)), and subsequently their harmonic average, the F1 score (2 * (`precision` * `recall`) / (`precision` + `recall`)), utilizing the number of true positives (TP), false positives (FP), and false negatives (FN). In order to assess whether an element in the reconstruction is a positive or negative, we selected a decision boundary which generally maximized total F1 scores across realizations of NMF with various rank *k.* We find that this is the case for a decision boundary of 0.35 (**Extended Data Fig. 3d**).

We considered values of *k* between 4 and 36. While the increase in total F1 scores as a function of *k* is monotonic, the relation is sublinear pointing to diminishing returns (**Extended Data Fig. 3e**). This effect seemed to become prevalent somewhere between 10 and 20 components. Calculating the derivative of the F1 score relative to the number of components *k*, showed that *k*=16 is a point where adding additional components leads to diminishing returns (**Extended Data Fig. 3f**). We therefore chose to use *k*=16 components for downstream analyses, for a relatively good reconstruction performance while allowing for a simplified lower-dimensional interpretation.

We used the Python scikit-learn package version 0.19.1 implementation of NMF. We used a coordinate descent solver for the Frobenius norm objective function. Non-negative double singular value decomposition (NNDSVD)[23] was used to initialize the weights. A random seed of 20 was used for the SVD algorithm. Other parameters were left as default: tol=0.0001, max_iter=200, alpha=0.0, l1_ratio=0.0. An implementation of our setup is available (https://github.com/Altius/DHSVocabulary).

##### Labeling of NMF components and DHSs

To aid interpretation of the 16 NMF-derived components, we used two orthogonal approaches to assign labels to components, based on (i) biosample properties and (ii) DHS sequence features.

First, for each component we obtained the top 15 DNase-seq datasets in terms of how much of their signal is attributed to that component. In other words, we selected biosamples based on component-specific NMF loadings present in their datasets (**Extended Data Fig. 4a**). These maximal NMF loadings across datasets were generally strong across components (**Extended Data Fig. 4b**). In general, a clear pattern emerged of shared properties of biosamples most strongly associated with specific components. To generalize this, we systematically performed one-sided Mann-Whitney U tests to assess whether NMF loadings for biosamples sharing certain metadata categories are greater than those for biosamples not in the given metadata category (**Extended Data Fig. 4c**). In particular, we assessed metadata categories corresponding to human organ systems and the cancer status of biosamples. P-values were corrected for multiple hypothesis testing using the Bonferroni correction method. A post-hoc analysis of biosample-to-component assignment for values of $k < 16$ provided insight into the genesis of our $k$=16 component model, showing junctures after which separate cell type lineages are captured by distinct components (**Extended Data Fig. 4d**).

Second, for each component we obtained DHSs with maximal NMF loadings for that component, and subsequently performed enrichment analyses for transcription factor (TF) binding site motifs (**Extended Data Fig. 5a**). We employed a wide array of TF motifs and use FIMO[25] (match threshold $p < 10^{-5}$), to search for motif instances in the human genome. We tested the association of motif occurrences with specific NMF components using Fisher's Exact test. We grouped similar motifs into clusters (https://resources.altius.org/~jvierstra/weblogos/main.html) for the purpose of summarization and visualization. The results show strong enrichments for component-specific motifs, corresponding to the putative binding of component-relevant transcription factors (**Extended Data Fig. 5b**).

Taken together, the strong associations of 1) biosample properties and 2) transcription factor binding site occurrences with specific components, enabled a labeling of each NMF component, resulting in a DHS Vocabulary (**Fig. 2d**). For downstream analysis, we labelled each DHS with its strongest NMF component (**Fig. 2e**). The DHS Index annotated with these strongest NMF Vocabulary components is available via the ENCODE DCC portal (https://www.encodeproject.org/annotations/ENCSR857UZV/) and via Zenodo (https://doi.org/10.5281/zenodo.3542126).

#### Regulatory annotation of human genes

##### Per-component genomic distribution of DHSs

Same-component DHSs tend to occur closer together than expected by random chance, as compared against empirical distributions based on random assignment of component labels to DHSs and sampling the same number of DHSs 1000 times (**Extended Data Fig. 6a**).

##### Per-component meta-DNase tracks

To illustrate regional diversity of DHS component data, we generated data tracks representing each of the 16 DHS component (**Extended Data Fig. 6b**), as follows. For each component, we obtained the top 15 datasets most strongly associated with the component (**Extended Data Fig. 4a**). We then calculated meta-DNase tracks for each component by averaging genome-wide DNase-seq signal profiles across these top 15 datasets. For visual conciseness, we also provide aggregate tracks that overlay the meta-DNase tracks of all Vocabulary components (as shown in e.g. **Fig. 3c, Extended Data Fig. 6c, Extended Data Fig. 7-9**).

##### Definition of regulatory landscapes and component associations

We defined the regulatory landscape of a gene as the set of DHSs within the gene body, plus DHSs in flanking regions of maximally 5kb upstream and maximally 1kb downstream of the gene body, or up until halfway through to the gene upstream, whichever value is smaller (**Fig. 3a**). This prevents flanking region DHSs from being routinely assigned to the regulatory landscapes of multiple genes and alleviates mixing of regulatory signals.

We tested the regulatory landscapes of all 56,832 annotated GENCODE genes (**Fig. 3b**) for association with DHS Vocabulary components for each component separately. Specifically, under the null hypothesis that DHS components are randomly distributed across gene regulatory landscapes, we used the binomial distribution to test the hypothesis that the proportion of DHSs annotated with a given component is higher among DHSs within a particular gene regulatory landscape than outside. We controlled the False Discovery Rate (FDR) at 5% by calculating q-values[26] across the total of all genes and components.

##### Annotations for GENCODE genes and pseudo-gene types

GENCODE v28 (Basic) annotations were used for all analyses. For the purposes of labeling and visualizing genes, we obtained for each gene the longest transcript as its representative region. Pseudo-gene annotations, including information on parent genes, were obtained from psiCube (http://www.pseudogenes.org/psicube/)[27]. In particular, we used the following annotation file: http://pseudogene.org/psicube/data/gencode.v10.pgene.parents.txt.

##### Visualization of gene regulatory annotations

We used t-Distributed Stochastic Neighbor Embedding (t-SNE) to visualize the enrichment ratios of gene regulatory landscapes, in terms of their association with DHS components (**Fig. 3d, Extended Data Fig. 9**). Each dot shown represents a gene found to be significantly associated with one or more DHS components, and the union of these are the genes used to calculate the 2D embedding. The R (http://www.r-project.org) implementation as provided in the `Rtsne` package was used, with default parameters. Genes are colored according to their (most strongly enriched) significant DHS component.

##### Construction and use of gene expression compendium dataset

We employed the full human ARCHS4 dataset (downloaded June 26, 2018)[28] and selected relevant tissue and cell types for each DHS Vocabulary component. Specifically, we used regular expressions to globally describe and filter on tissue and cell types in the available ARCHS4 dataset metadata, as follows:

Placenta: `placenta|trophoblast`
Lymphoid: `lymph|T-cell|B-cell|CD[48]|regulatory|thymus`
Myeloid/erythroid: `myel|CD34|erythr|cord blood|KBM7|K562`
Cardiac: `cardiac|cardio|heart|ventricle`
Musculoskeletal: `muscle|bone|skeletal|myocyte|tongue EXCEPT marrow|cardio|cardiac`
Vascular/endothelial: `vascul|endotheli|vessel|vein|arter|HMVEC EXCEPT H1`
Embryonic/Primitive: `embryonic|iPSC|H1|H9 EXCEPT derived|fibroblast`
Neural: `neuro|nervous|brain|glia|eye|retina|spinal EXCEPT H1`
Digestive: `digestive|colon|liver|intestin|hepato`

Stromal A & B: `fibroblast EXCEPT iPSC|cardiac`
Renal/cancer: `renal|kidney|podocyte|glomeru|tubular EXCEPT fetal`
Cancer/epithelial: `epithelial|keratino|melanocyt|MCF`
Pulmonary devel.: `fetal AND lung EXCEPT fibroblast`
Renal/organ devel.: `fetal AND kidney`

Datasets occurring in multiple groups, with the exception of Stromal A & B, were removed to prevent ambiguity. This resulted in a total of 33,733 unique gene expression datasets, with expression information for 35,238 genes.

For each of the reported 35k+ genes, we obtained the 95th percentile value across datasets selected for each Vocabulary component as the representative value for that gene in that component. We chose the 95th percentile to not be led by outliers in the data, while still being sensitive for cell type selective expression levels. For each DHS component, we calculated average expression levels across genes labeled with that component ("observed"), as well as across all component-labelled genes ("expected"). The resulting values are reported as $\log_2$-transformed observed/expected ratios (**Fig. 3g**).

##### Annotation and visualization of pathway labellings

A curated set of canonical pathways was obtained from the MSigDB Collections (http://software.broadinstitute.org/gsea/msigdb/genesets.jsp?collection=CP). Pathway enrichment analyses (**Extended Data Fig. 10**) were performed analogously to gene enrichment analyses, by pooling DHSs in regulatory neighborhoods of all pathway-associated genes. We employed the KEGG REST API (https://www.kegg.jp/kegg/rest/keggapi.html) to download and graphically annotate KEGG pathway representations.

#### Genetic variation analyses

##### GWAS traits and summary statistics

We obtained Genome-Wide Association Study (GWAS) summary statistics data from the UK Biobank project as processed by the Neale lab (http://www.nealelab.is/uk-biobank/). Additionally, we obtained GWAS summary statistics calculated using BOLT-LMM v2.3[29], as used in recent work[30].

##### Estimates of SNP-based heritability

GWAS traits were curated by removing those with a narrow-sense SNP-based heritability ($h_g^2$)[31] of less than 1%. Although ideally we would quantify heritability by considering the true causal effects of variants, in reality we do not observe these. Instead, we are limited to GWAS summary statistics, which essentially describe the marginal trait-correlation for each variant, consisting of both causal effects and effects due to Linkage Disequilibrium (LD), plus statistical noise. Recently proposed methods such as LD score regression (LDSC)[32] are able to estimate heritability while explicitly considering the underlying LD structure. For continuous traits, in case both raw and inverse-ranked normalized (irnt) versions were available, we retained the latter only. This yielded a total of 1,316 traits for subsequent analyses with an $h_g^2$ of at least 1%.

**Quantitative trait associations**
For quantitative trait-versus-component analyses (**Fig. 4a, Extended Data Fig. 11**), we assessed the correspondence between trait association strength (GWAS variant association p-value) and the component annotations of variant-harboring DHSs, for increasingly stringent subsets of GWAS variants. Enrichment p-values were calculated using a binomial distribution, as done previously[33]. We explicitly control for large scale Linkage Disequilibrium (LD) structure, using a form of LD clumping[34], by selecting a single variant-containing DHS for each of 1,708 approximately independent LD blocks[35]. Namely, for each LD block, the variant with the lowest GWAS association p-value that overlaps a DHS was selected for subsequent analysis. In case multiple such variant-containing DHSs exist, we gave preference to the DHS with the highest confidence score (mean signal) in our DHS Index.

**Stratified LD-score regression**
To estimate $h_g^2$ with maximal statistical power, we employed LD score regression (LDSC)[32] to explicitly take into account LD structure. In particular, we employed a stratified version of LDSC (S-LDSC)[36] to partition heritability estimates according to pre-defined sets of genome-wide annotations (**Fig. 4b-c, Extended Data Fig. 12**), consisting of our annotated DHSs in addition to a wide range of 85 genome-wide functional 'baseline' annotations (baseline-LD model v2.1). The v2.1 baseline set consists of a total of 86 genome-wide annotations, building upon the 76 annotations used in the v2.0 set and several additional annotations[37]. These 'baseline' annotations encode whether SNPs fall inside protein-coding or non-coding regions, regions with increased levels of evolutionary conservation, regions predicted or confirmed to have enhancer activity, etcetera. Their breadth provides a robust[36] baseline model along which to test trait heritability contributions of our DHS components. We express the heritability enrichment of an annotation as the ratio of its proportion of per-trait $h_g^2$ and the proportion of SNPs covered by the annotation (**Fig. 4b**).

Variants included in the analysis are those registered in HapMap3, with a minimal minor allele frequency (MAF) of 5%, and excluding the human Major Histocompatibility Complex (MHC) locus. Baseline LD scores were computed from 1000 Genomes Phase 3 data from European ancestry populations and corresponding allele frequencies (as used previously[37] and available from the LDSC reference downloads page, along with the baselineLD annotation set — https://data.broadinstitute.org/alkesgroup/LDSCORE/).

**Heritability enrichments for DHS Vocabulary components**
We applied S-LDSC to our DHS Vocabulary components as follows. Briefly, each DHS was assigned to its majority DHS component and (when possible) assigned to overlapping variants. For the resulting Vocabulary-based annotations, LD-scores were calculated. We then performed S-LDSC separately for each of the selected 1,316 traits, relative to these Vocabulary-based annotations and the baselineLD model described above. For each trait vs. annotation combination, we obtained estimates of its heritability enrichment, expressed as the ratio of its proportion of $h_g^2$ and the proportion of SNPs covered by the annotation (**Fig. 4b-c**). We considered heritability enrichments statistically significant at an estimated False Discovery Rate (FDR) of less than 5% calculated across all considered traits and DHS components. Note that this is more stringent than the commonly used per-trait correction for multiple hypothesis testing.

##### Unique per-annotation contributions to SNP-based heritability

Estimates of heritability enrichment can be confounded by contributions of multiple (overlapping) genomic annotations included in S-LDSC models. To quantify unique per-annotation contributions to heritability, we obtained the average per-SNP increase in heritability ascribed to that component, after controlling for all other annotations in the model (baseline annotations and DHS components). We report derived Z-scores for each trait-versus-component combination (**Fig. 4d**).

To quantify the heritability contribution of per-dataset DHSs, we performed a variation on the standard S-LDSC procedure, as described previously[36]. Specifically, we built upon the baselineLD model, by iteratively considering annotations derived from individual datasets only. These individual datasets were collected by selecting for each trait the 15 datasets most informative to each DHS component (**Extended Data Fig. 4a**). Annotations consist of DHSs observed in those datasets, as well as their complement, i.e. the remainder of Index DHSs. We report the contribution to heritability based on the former, expressed as Z-scores (**Fig. 4d-e**).

### Data and Code Availability

#### Biosample metadata

Biosample metadata are available in Supplementary Table 1 as well as in a variety of other formats via Zenodo (https://doi.org/10.5281/zenodo.3542126).

#### Index of consensus DHSs

Code used for constructing the index of consensus DHSs is available via Github (https://github.com/Altius/Index).

The resulting set of 3.5M+ DHS delineations is available in tab-separated format from the ENCODE DCC portal (https://www.encodeproject.org/annotations/ENCSR857UZV/) and via Zenodo (https://doi.org/10.5281/zenodo.3542126).

#### Construction of DHS Vocabulary

Data matrices describing the occurrence patterns of DHSs across biosamples are available in various formats via Zenodo (https://doi.org/10.5281/zenodo.3542126).

Code used for the decomposition of DHS occurrence patterns across biosamples is available via Github (https://github.com/Altius/DHSVocabulary).
